# Stochastic colonization and host-to-host transmission shape gut bacterial variability

**DOI:** 10.64898/2026.05.11.724410

**Authors:** Carol Lu, Stanimir Asenov Tashev, Pedro Pessoa, Rory Kruithoff, Douglas P Shepherd, Steve Presse

## Abstract

Understanding the kinetic processes that govern bacterial population dynamics within hosts is critical in developing effective strategies to control microbiota. However, inferring population dynamics is challenging due to large host-to-host bacterial population variability stemming from stochastic colonization events, as well as the inability to continuously monitor the bacterial population without disturbing the host. Using *C. elegans* fed *E. coli* under different diets, we show that early colonization acts as a stochastic bottleneck that drives substantial divergence in host-level bacterial loads, and that the spreading of bacteria from colonized worms to sterile ones regulates this variability by altering effective colonization pressure. These conclusions are reinforced using a simulation-based inference framework that quantifies stochastic within-host population dynamics from discrete snapshot data, enabling inference of effective colonization and growth rates across heterogeneous hosts with variable carrying capacities. Applying this framework, we further demonstrate that the bacterial predator *B. bacteriovorus* reduces average gut bacterial loads by two orders of magnitude, primarily by suppressing environmental recolonization and subsequent host-to-host transmission rather than eliminating established intra-host populations. Together, these results reveal that host-associated microbial population dynamics are strongly impacted by environmental colonization processes that modulate stochastic entry events.

## 1 Introduction

The gut microbiome plays a central role in animal health, influencing metabolism, immunity, and disease susceptibility [1] and its dynamics have been studied across a range of model organisms, including zebrafish [2, 3] and other vertebrate and invertebrate systems [4–6].

Disruptions to this ecosystem (dysbiosis) are linked to a wide range of pathological conditions [7, 8]. Despite extensive characterization of microbial composition [9], a quantitative understanding of the dynamic processes governing microbial populations *in vivo* remains elusive. This is, in part, due to challenges in quantifying microbiome kinetics from direct observation, which hinders our ability to predict microbial responses to interventions, such as antibiotic treatment, an urgent challenge in the face of rising antibiotic resistance [10, 11].

Toward understanding gut microbiome dynamics, our focus here is on gut infection in the nematode *Caenorhabditis elegans* (*C. elegans*), a well-established model for host–microbe interactions [12]. *C. elegans* benefits from a highly characterized physiology and genetic toolkit [13–16] and is known to share susceptibility to a variety of pathogens also affecting mammals, including humans [17, 18], such as *Escherichia coli* (*E. coli*). Although *E. coli* is widely used as a food source for *C. elegans* [13, 19, 20], it can exert deleterious effects on its host [21, 22]. In particular, intestinal accumulation of *E. coli* can impair host physiology, including reduced lifespan and intestinal dysfunction [22, 23], making *C. elegans* and *E. coli* together a rich setting in which to study infection dynamics and microbial population behavior *in vivo*.

While previous studies of gut microbiota dynamics have sought to link bacterial composition to host health using measurements within individual hosts [24–26], dysbiosis and related pathologies are often initiated by rare infection from the environment. Such infection events trigger gut bacterial population growth [6, 27, 28] and alter gut microbiome stability [29]. However, the underlying colonization and growth processes cannot be uniquely inferred from host-level bacterial load alone. Consequently, quantitative kinetic models of not only in-host bacterial growth but also environmental effects and potential host-to-host transmission are required in describing the eventual microbial distributions across hosts.

The challenge in kinetic modeling, however, lies in the fact that stochastic colonization events at short timescales and large populations within hosts at longer timescales impose upon us a fundamentally multiscale modeling problem [20, 29]. Estimating growth parameters across this multiscale regime by averaging bacterial loads across hosts and fitting deterministic models such as logistic growth is inappropriate because such fitting assumes host-to-host variability reflects fluctuations around a shared trajectory [30]. However, the shared trajectory assumption is problematic in describing the substantial bacterial load heterogeneity between individuals [31] because it ignores small initial number effects derived from infection. Capturing these initial effects, therefore, requires measurements capable of resolving viable bacterial populations across several orders of magnitude, including very low-abundance colonization states.

Although specialized imaging and flow-sorting approaches can achieve single-cell resolution, common methods such as fluorescence imaging [20], qPCR [32], and bulk growth assays [33] often provide indirect or relative abundance measurements rather than absolute viable bacterial counts. We therefore quantified gut-associated bacterial populations using dilution plating of crushed worms to obtain colony-forming unit (CFU) measurements across hosts and time points. Yet quantitatively inferring the kinetic parameters governing bacterial population growth remains challenging because bacterial counts are measured as sparse, discrete snapshots rather than continuous trajectories. Therefore, we combined probabilistic reconstruction of bacterial populations from colony counts using our REPOP (REconstruct POPulations from plates) framework [34] with a hybrid simulation-based inference framework [35–37] that bridges stochastic low-copy-number dynamics [38–40] and continuum descriptions of large populations [41–43]. To experimentally separate colonization from downstream growth, we further designed feeding conditions that systematically varied the timing and magnitude of bacterial exposure.

Through our combined experimental and modeling framework, we disentangle population-level gut dynamics across scales, both within hosts and induced by environmental factors. By modifying the diet of the nematodes to change kinetic rates, we show that stochastic early colonization strongly determines eventual bacterial loads, while environmental exposure modulates both bacterial load and host-to-host variability, implicating ongoing recolonization and transmission between hosts through the environment. Applying this framework to perturbations with the bacterial predator *Bdellovibrio bacteriovorus* (Bb), we further find that reductions in gut bacterial load arise primarily from suppressed environmental recolonization between hosts rather than enhanced clearance of established gut populations.

## 2 Results

To understand the interplay between stochastic colonization and reseeding/transmission, we designed a set of experiments systematically varying the timing and magnitude of bacterial exposure. These perturbations were designed to test how early stochastic colonization introduces host-to-host variability, and assess whether population dynamics are governed solely by within-host processes or strongly influenced by environmental reseeding and transmission. To further probe the role of environmental coupling on both colonization and reseeding/transmission, we introduced the bacterial predator Bb as a targeted perturbation that can suppress environmental bacterial abundance while also potentially acting within the host.

To quantitatively capture bacterial populations spanning early colonization to later large-population expansion, we employed serial dilution and plating, as highlighted in Fig. 1a, to enumerate CFUs from viable *E. coli* that remained associated with worms after removing external bacteria through washing, purging, and surface bleaching. These CFU measurements were therefore taken as estimates of gut-adhered bacterial populations. This approach yields absolute population sizes necessary for quantitative modeling of the kinetic rates governing *E. coli* population growth, including colonization, replication, loss, and host capacity (Fig. 1b).

**Figure 1:**
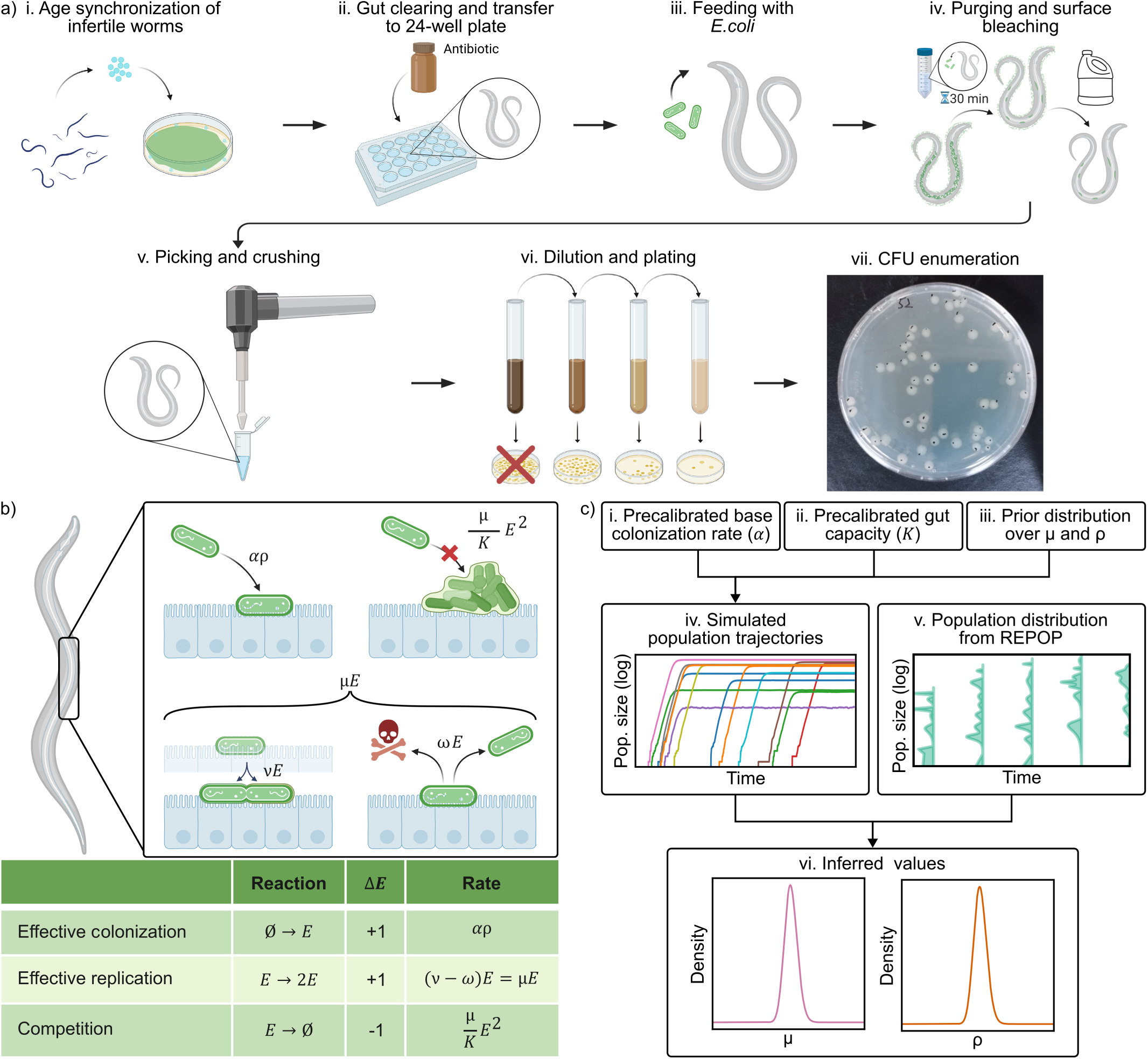
Bacterial population dynamics in *C. elegans* alongside our experimental and computational pipeline. a) To obtain quantitative bacterial counts from individual worms, we follow this experimental protocol, detailed in Sec. 4.1: i. Eggs are harvested from mature worms and then grown for 2 days at a non-permissive temperature to generate infertile adults. ii. Worms are moved to a 24-well plate and incubated with heat-killed bacteria and an antibiotic for a day to clear the gut of the worms. iii. Worms are fed different diets for a period of time, as detailed in Sec. 4.1. iv. Every 48 h worms are collected from a well, washed, purged by incubating with heat-killed bacteria, and surface-bleached. v. Individual worms are picked and homogenized. vi. The homogenates are diluted and then plated. vii. After overnight incubation, the CFUs are counted. b) Graphical representation of the biological mechanisms we want to quantify: colonization, density-dependent competition, replication, death, and expulsion. The effective rate of colonization depends on the maximal colonization rate *α*, when the worms are fed on a regular live *E. coli* diet (see Fig. 2), scaled by the environmental parameter *ρ*, which depends on the colonization-competent bacteria in the environment. The death and expulsion rates are combined in *ω*, which is then subtracted from the total replication rate *ν* to give the effective rate of replication *µ*. The competition between bacteria inside the gut leads to a maximum worm capacity *K*. c) To infer the values of *ρ* and *µ*, we first precalibrated *α* and *K* (i. and ii.) from experimental results, while *ρ* and *µ* parameters are drawn from priors (iii.). We simulate population trajectories from these parameters using a multiscale framework (iv.) and combine the resulting population distribution with the population distributions we reconstruct from CFU counts using REPOP [34] (v.) to calculate the posterior probability distributions over *ρ* and *µ* (vi).

Finally, to extract key parameters from our multi-scale data spanning individual bacterial colonization to large-scale steady-state populations, we developed a simulation-based inference approach quantifying the effects of colonization and reseeding/transmission from snapshot data absent continuous observation (Fig. 1c).

### 2.1 Impact of magnitude and timing of live bacterial feeding on gut microbial count

To disentangle the contributions of early stochastic colonization and continued environmental recolonization to gut bacterial population dynamics, we designed feeding conditions that systematically varied both the magnitude and duration of exposure to live *E. coli* (Fig. 2a). Bacterial loads within individual *C. elegans* guts were quantified using serial dilution plating adapted from Ref. [20], followed by probabilistic reconstruction of host-level bacterial load distributions using REPOP [34].

**Figure 2:**
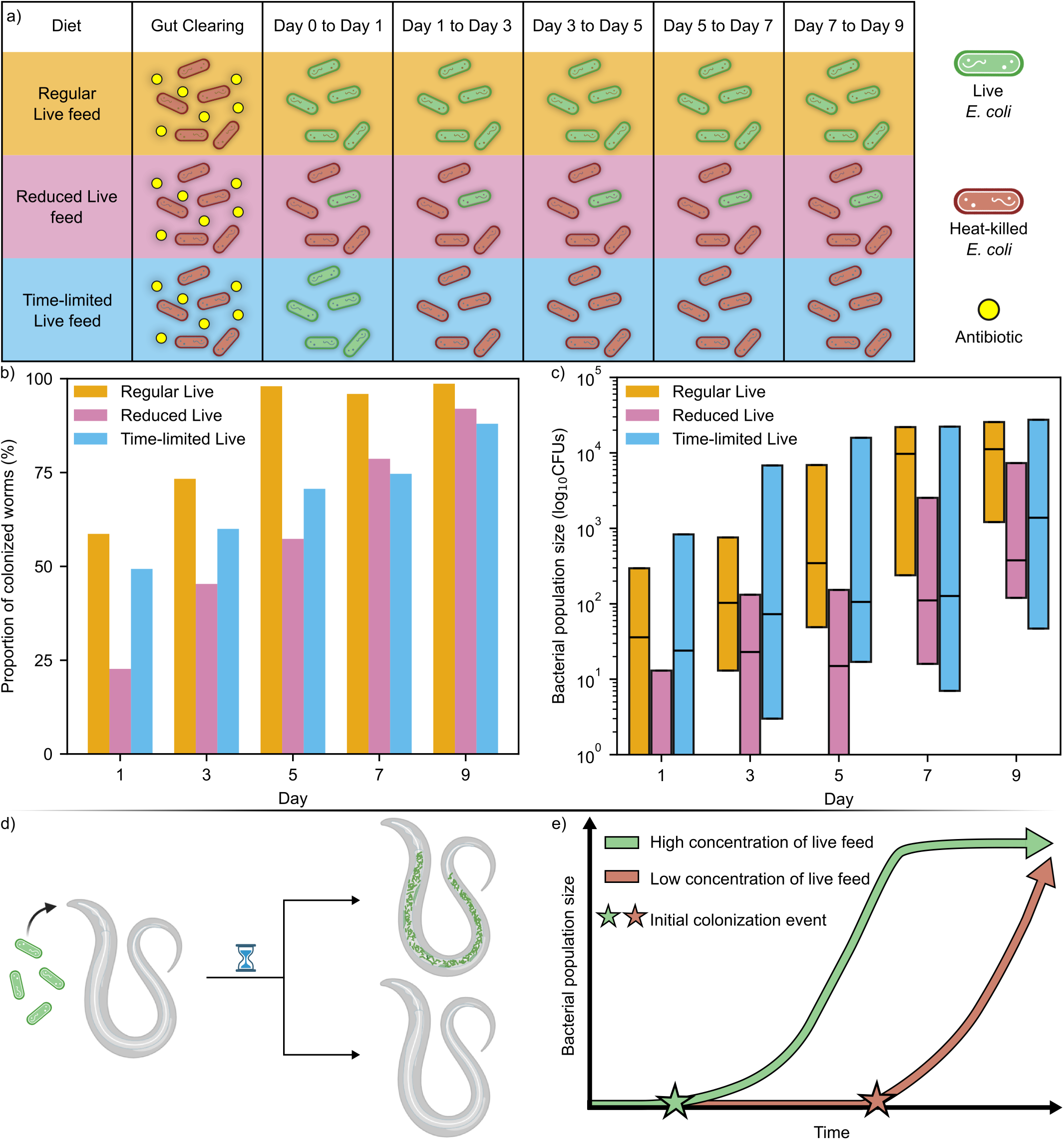
Timing and magnitude of live bacterial exposure shape gut bacterial load and variability. a) Worm diets featuring different periods of live (green) and heat-killed (red) bacteria, after an initial 24 h of gut clearing with an antibiotic (yellow). These include: a continuous high-concentration live *E. coli* (regular live), a constant 100-fold reduced live concentration (reduced live), or full live exposure for 24 h followed by heat-killed bacteria (time-limited live). b) Mean percentage (across 3 repeats with combined n = 75 worms per diet) of colonized worms from the three diets. Regular live feeding (orange) yielded almost full colonization by day 3, while reduced live feeding (pink) delayed colonization. Despite the removal of live bacteria after day 1, the fraction of colonized worms in the time-limited (blue) condition continued to increase after. c) Box plots showing reconstructed quartiles of gut bacterial population sizes under the regular live (orange), reduced live (pink), and time-limited (blue) diets. Regular live feeding produced increasing bacterial loads approaching a plateau at later time points, whereas reduced live feeding maintained substantially lower population sizes. In contrast, the time-limited live diet exhibited broadened variability at later time points. d) Cartoon showing the stochasticity of colonization after incubation with live bacterial feed. e) An illustration showing how the time of initial colonization events (stars) in diets with low concentration (orange) of live *E. coli* can lead to a delayed progression of the gut microbiota numbers, compared to higher live bacterial concentrations (green).

Age-synchronized bacteria-free adult worms were established in 24-well plates and maintained on diets in which live *E. coli* were partially or temporally replaced with heat-killed bacteria. This substitution preserved overall bacterial biomass exposure while selectively modulating the availability of colonization-competent cells. At each feeding interval, individual worms were isolated and plated to quantify viable gut-associated bacteria (Fig. 1a).

Three feeding regimes were designed to probe distinct colonization dynamics. Under the regular live diet (first row of Fig. 2a), worms were continuously exposed to high concentrations of live *E. coli* throughout the experiment, representing sustained high colonization pressure. Under the reduced live diet (second row of Fig. 2a), worms received a 100-fold lower concentration of live bacteria supplemented with heat-killed bacteria, representing persistent but weak colonization pressure. Finally, under the time-limited live diet (bottom row of Fig. 2a), worms were exposed to live *E. coli* only for the initial 24 hours before being washed to remove surface-associated live bacteria and transferred to heat-killed *E. coli*, representing a transient period of high colonization pressure.

Through the regular live diet, we first establish that colonization of the worm gut arises from stochastic establishment events, not guaranteed to occur in every host. More specifically, despite sustained high exposure, colonization remained incomplete at early time points: after 24 h, 59% of worms were colonized, with this fraction increasing over time to nearly complete colonization by day 5 (98%) (Fig. 2b and Tab. S1). Also, consistent with stochastic colonization, bacterial loads varied substantially across individual worms at each time point within the regular live diet. Using REPOP reconstructions, we found that under the regular live diet, bacterial loads spanned a broad range at early time points, with a median of 36 CFUs per worm [0–296 CFUs] at day 1 (Fig. 2c and Tab. S2). Here, square brackets denote the interquartile range (IQR), that is, the 25th to the 75th percentile, reconstructed from the inferred bacterial load distributions. Bacterial loads increased over time and approached a plateau at later time points, reaching 9705 CFUs [239–21960 CFUs] at day 7 and 11190 CFUs [1211–25609 CFUs] at day 9. The plateau on the order of 10^5^ was consistent with previous estimates of *C. elegans* gut capacity [20].

In contrast, the reduced live diet predictably yielded substantially lower bacterial loads across all days (Fig. 2c). On day 1, the 75th percentile bacterial load was 13 CFUs, nearly an order of magnitude lower than in the regular live diet. Moreover, by day 9, the median bacterial load only reached 377 CFUs [120–7330 CFUs] (Tab. S1). While bacterial loads increased over time, the worm population as a whole did not approach the high-bacterial-population plateau observed under regular feeding (illustrated alternatively in Fig. S1a).

In the time-limited live diet, early time points were similar to the regular live diet, with 49% of worms colonized at day 1 and a median bacterial load of 24 CFUs [0–832 CFUs]. However, following the transition to heat-killed feeding by hour 24, the distribution of bacterial loads diverged from the regular live diet. Although the inferred 25% of the most highly colonized worms still reached bacterial loads comparable to those under the regular live diet, around 20000, the distribution broadened substantially at later time points, with the inferred 25th percentile remaining at fewer than 20 CFUs even through day 7 (Tab. S2). This broadened distribution was consistent with stochastic initial worm colonization, in which later populations comprise both highly colonized and weakly colonized worms. However, the effect of the broadened distribution decreased over time, with the final time-limited live diet worms’ inferred 25th percentile rising to 47 CFUs (Tab. S2). Rather than maintaining the expected proportion of non-colonized worms, the fraction of colonized worms continued to increase over time (Fig. 2b), suggesting an environmental reseeding component (Fig. S2a).

To assess whether the stochastic establishment events and broadened bacterial load distributions observed under time-limited live diet simply a consequence of the original media conditions, we repeated the regular live and time-limited live diets in modified S medium. Specifically, the standard S medium was adjusted from pH 5.8 to pH 7.3 (Fig. S3a), which is closer to the near-neutral conditions that are more favorable for *E. coli* growth [44]. Under these conditions, the fraction of colonized worms at day 1 remained similar to that observed at pH 5.8, with 63% and 56% colonization in the regular live and time-limited live diets, respectively (Fig. S3b and Tab. S3). Similarly, median gut bacterial loads at day 1 were comparable between the two diets (21 and 32 CFUs for regular live and time-limited live diet, respectively). At later time points, however, the time-limited live diet at pH 7.3 diverged from earlier results at pH 5.8, with lower bacterial population sizes relative to the regular live diet (Fig. S3c and Tab. S4). Nevertheless, consistent with observations at pH 5.8, the fraction of colonized worms in the time-limited live diet increased over the course of the experiment (Fig. S3b), again indicating a potential recolonization effect. Inspecting the environment in the time-limited feeding experiment, we observed a significant amount of live *E. coli* on days 3 to 9 (Fig. S4a). To reduce the abundance of live bacteria in the environment and magnify the effect of stochastic early colonization, we next turned to using bacterial predators.

### 2.2 Effects of predation on worm gut microbiota

To investigate environmental reseeding and transmission without suppression using chemical antibiotics, we introduced a targeted perturbation to the system using the gram-negative bacterial predator Bb [45, 46]. This perturbation is of particular interest because prior studies have shown that Bb can reduce gut bacterial loads in *C. elegans* while also extending host lifespan [47], suggesting a potential role as a non-chemical antimicrobial agent.

Initial experiments revealed that the low pH (5.8) and high salinity (100 mM) of the original S medium did not support Bb predation (Fig. S5a,b), consistent with previous studies suggesting that Bb activity is inhibited under high osmolarity and low pH conditions [48–50]. Consistent with this, even under pH-adjusted (7.3) conditions at high salinity, we observed only minimal changes in percent colonization (Fig. S5b, Tab. S3), population distributions (Fig. S3c, Tab. S4), and environmental bacterial levels (Fig. S3, Fig. S4b), suggesting limited predation activity. To enable effective predation, we reformulated the medium to reduce the salinity tenfold prior to adjusting the pH to 7.3, and observed significant lysis of *E. coli* (Fig. S5c).

In order to establish whether the number of colonized worms and bacterial load distributions change over time under predation, we studied the effects of Bb predation using bacteria-free worms exposed to the time-limited live diet in the low salinity, pH 7.3 S media. This feeding regime was specifically chosen to isolate the effects of Bb on bacteria already associated with the worms, rather than on continuously replenished environmental *E. coli*. The control condition (without Bb) and Bb-treated condition are shown in the top and bottom rows of Fig. 3a, respectively. Both control and predator-exposed worms received a single 24 h exposure to full-concentration live bacteria followed by heat-killed feeding only. On heat-killed feeding days, the predator-exposed group was supplemented with Bb at OD_600_ = 0.2 per milliliter.

**Figure 3:**
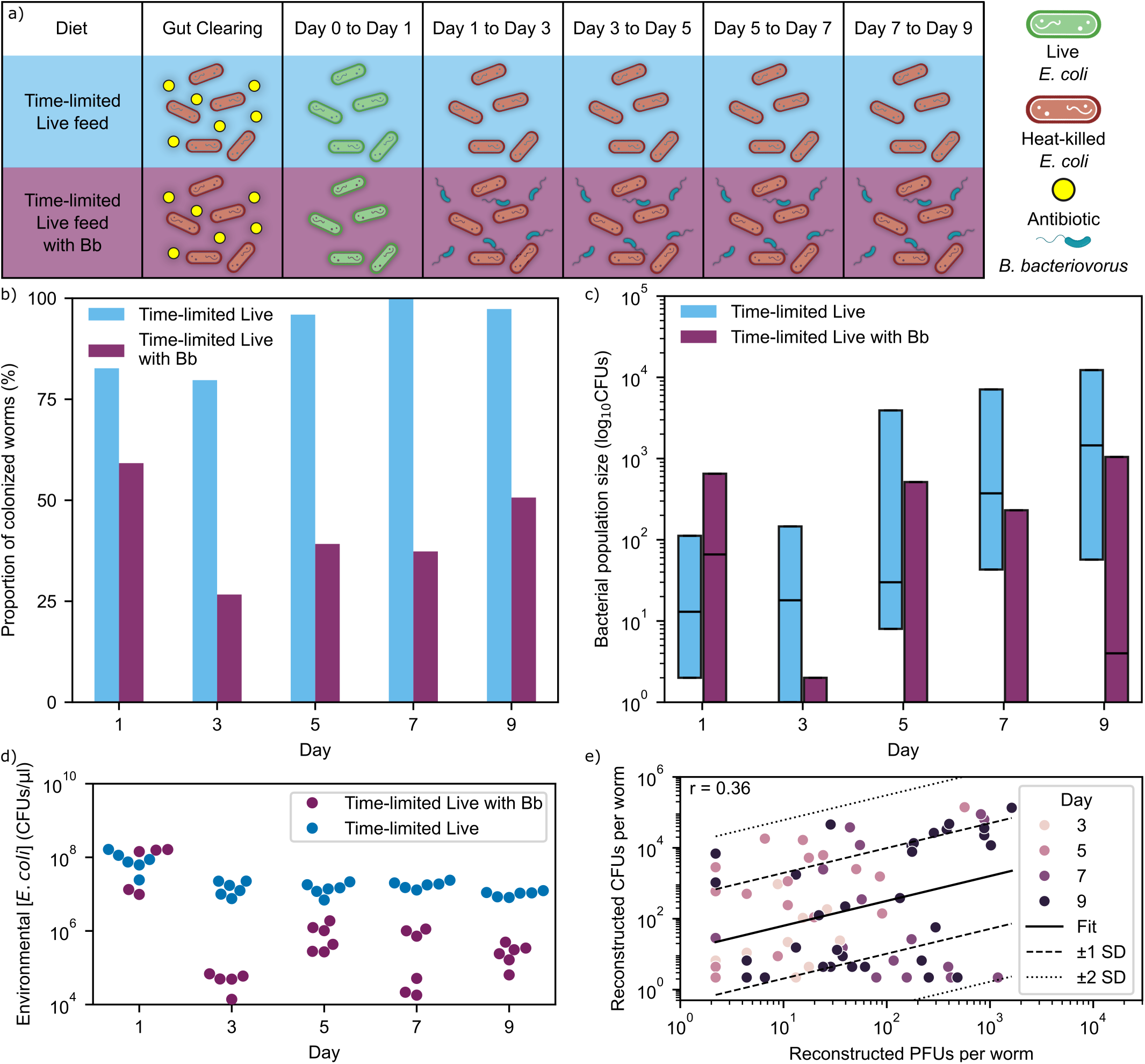
Predation depletes environmental bacteria and reduces the proportion of colonized worms, yet predator and prey remain uncorrelated within worm guts. a) Feeding schedule for the control and Bb-treated conditions. Worms were first cleared of gut bacteria with an antibiotic (yellow), exposed to live *E. coli* (green) for 24 h, and then maintained on heat-killed bacteria (red). Bb-treated worms additionally received Bb (dark blue) supplementation during the heat-killed feeding phase. b) Mean percentage of colonized worms with and without Bb treatment in purple and blue, respectively. Colonization remained persistently lower in the presence of Bb throughout the experiment. Day labels represent the time point at which worms were sampled. Sampled worms were not re-fed with bacteria on the day of crushing. c) Box plots showing reconstructed quartiles of bacterial loads per worm for the control and Bb-treated conditions in purple and blue, respectively. Bb treatment shifted bacterial load distributions toward lower values, although a subset of worms still reached high bacterial loads at later time points. n = 75 worms per diet across 3 repeats per condition, except for the first day of the Bb predation experiment, where a single repeat was omitted due to uneven dilution (see Sec. 4.1). d) Reconstructed environmental bacterial counts at different time points inside the wells of the time-limited feed with Bb (purple) and without (blue) in low-salinity S medium. e) Distribution of CFUs and PFUs for each worm, with worms color-coded by day. Worms not colonized with both *E. coli* and Bb were excluded. Linear regression and standard deviation (SD) are plotted, with the r value shown on the plot.

Since we expected Bb predation to produce a larger fraction of worms with lower bacterial loads approaching the detection threshold, we increased the sensitivity of the plating assay by lowering the initial plating dilution tenfold. We did so in order to better distinguish truly uncolonized worms from hosts with a low gut–adhered population (see Sec. 4.1).

Under control conditions, the fraction of colonized worms increased from 82% of worms on day 1 to 96% by day 7 (Fig. 3b and Tab. S5). In contrast, in the presence of Bb, the proportion of colonized worms remained consistently low throughout the experiment, ranging from 59% at day 1 to 51% at day 9.

Consistent with the reduced fraction of colonized worms, the distribution of bacterial loads across worms was shifted toward lower values in the Bb-treated group compared to the control after day 1 (Fig. 3c). Unlike the control experiment, in which the proportion of highly colonized worms increased over time, the composition of bacterial loads in the Bb-treated group remained relatively stable during the later days of the experiment (Fig. S1b). Notably, however, a subset of worms still reached high bacterial loads at later time points, with the inferred 75th percentile bacterial load reaching 1047 CFUs by day 9 (Tab. S6). These observations now raised the question as to whether Bb reduces bacterial load primarily by limiting recolonization from the environment or by eliminating bacteria within already-colonized hosts.

To test whether Bb reduces the abundance of colonization-competent bacteria within the environment, we measured the abundance of live *E. coli* in the surrounding medium over time. We found that, in the presence of Bb, environmental *E. coli* concentrations were reduced by approximately 2–3 orders of magnitude compared to the control condition (Fig. 3d), indicating strong suppression of the external bacterial reservoir. We next asked whether Bb acts within the host to eliminate established bacterial populations. To assess whether Bb predation occurs within hosts, we quantified viable Bb plaque-forming units (PFUs) from individual worm homogenates using double-layer plating (see Sec. 4). This allowed us to measure both *E. coli* CFUs and Bb PFUs within the same worms. We observed only weak correlation between CFU and PFU counts (r = 0.36) (Fig. 3e), and no clear relationship between high Bb burden and reduced *E. coli* load across worms (Fig. S6). Together, these results suggest that Bb-mediated reduction of gut bacterial loads is not primarily driven by direct elimination of established within-host populations.

Importantly, because worms in the time-limited feeding regime were exposed to live *E. coli* only during the initial 24 h period and then washed, viable bacteria detected in the environment at later time points must arise from bacteria previously associated with colonized worms. Thus, environmental recolonization in this regime reflects transmission between hosts mediated by expelled bacteria (Fig. S2a). Together with the weak association between Bb burden and within-host *E. coli* abundance, these observations suggest that Bb primarily suppresses bacterial populations by reducing environmental reseeding and inter-host transmission rather than by directly increasing the death rate of *E. coli* inside the gut (Fig. S2b). We next tested this hypothesis quantitatively using our inference framework.

### 2.3 Simulation-based inference of bacterial population rates

While host-level bacterial load distributions reveal how stochasticity and predation reshape population outcomes, they do not directly identify the degree to which reseeding and transmission of *E. coli* from host to host is impacted by the presence of predators in the environment.

In particular, shifts in *E. coli* load distributions could arise from changes in colonization, replication, loss, or combinations thereof. For this reason, we infer the kinetic parameters governing bacterial population dynamics under all 5 experimental conditions shown in Fig. 2a and Fig. 3a.

The inference of kinetic parameters is complicated by the multiscale nature of bacterial population dynamics. Early colonization is governed by rare stochastic events involving only a few bacteria, whereas established populations can reach hundreds of thousands of cells. While low-copy-number dynamics are well described by stochastic simulation and master equation approaches [38–40], and large populations by continuum diffusion-like approximations [41–43], neither framework alone can efficiently capture both regimes within a unified description.

Since the probability of observing the measured bacterial load distributions cannot be computed directly across both stochastic and large-population regimes, we developed a hybrid simulation-based inference framework. Using shared kinetic parameters, the framework simulates bacterial population trajectories within individual worms across both regimes (Fig. 1b,c; see Sec. 4.2 for details). This work made use of a large number of stochastic simulations, which were parallelized for computational efficiency and executed on multiple GPUs.

To separate intrinsic colonization from environmental effects, we parameterized the effective colonization rate as *αρ*, where *α* represents the maximal colonization rate under fully viable feeding conditions and the scaling factor *ρ* captures environmental modulation introduced by the presence of colonization-competent *E. coli*(see Sec. 4.2 for more detail). In other words, across all 5 experimental conditions (Fig. 2a and Fig. 3a), the inferred colonization scaling factor *ρ* reflected the availability of live bacteria in the environment.

As our primary objective was to learn about the modulation of environmental reseeding, we independently precalibrated: 1) the maximal colonization rate, *α*, from binary colonization experiments that measured the probability of successful establishment as a function of the time under full bacterial exposure (see Sec. 4.1); 2) the carrying capacity, *K*, as a worm-specific parameter drawn from the final-day bacterial load distributions under the regular live diet (Fig. S7). These precalibrated quantities were then held fixed during subsequent inference of the remaining kinetic parameters. Additional details of the precalibration procedure are provided in Section 4.2.

We found *α* = 0.052 h^−1^ [0.045–0.060] for standard S medium (Fig. 4a), consistent with previous estimates [20]. Under full live feeding, we fixed *ρ* = 1 to represent saturating exposure to live bacteria, such that the rate of bacterial colonization is not limited by bacterial availability. For all other conditions, we estimated the parameter values *ρ*. We also inferred *µ*, which is the effective replication rate that combines the net effect of bacterial replication and loss (Fig. 1b) in the gut, as the latter cannot be independently quantified due to parameter indeterminacy.

**Figure 4:**
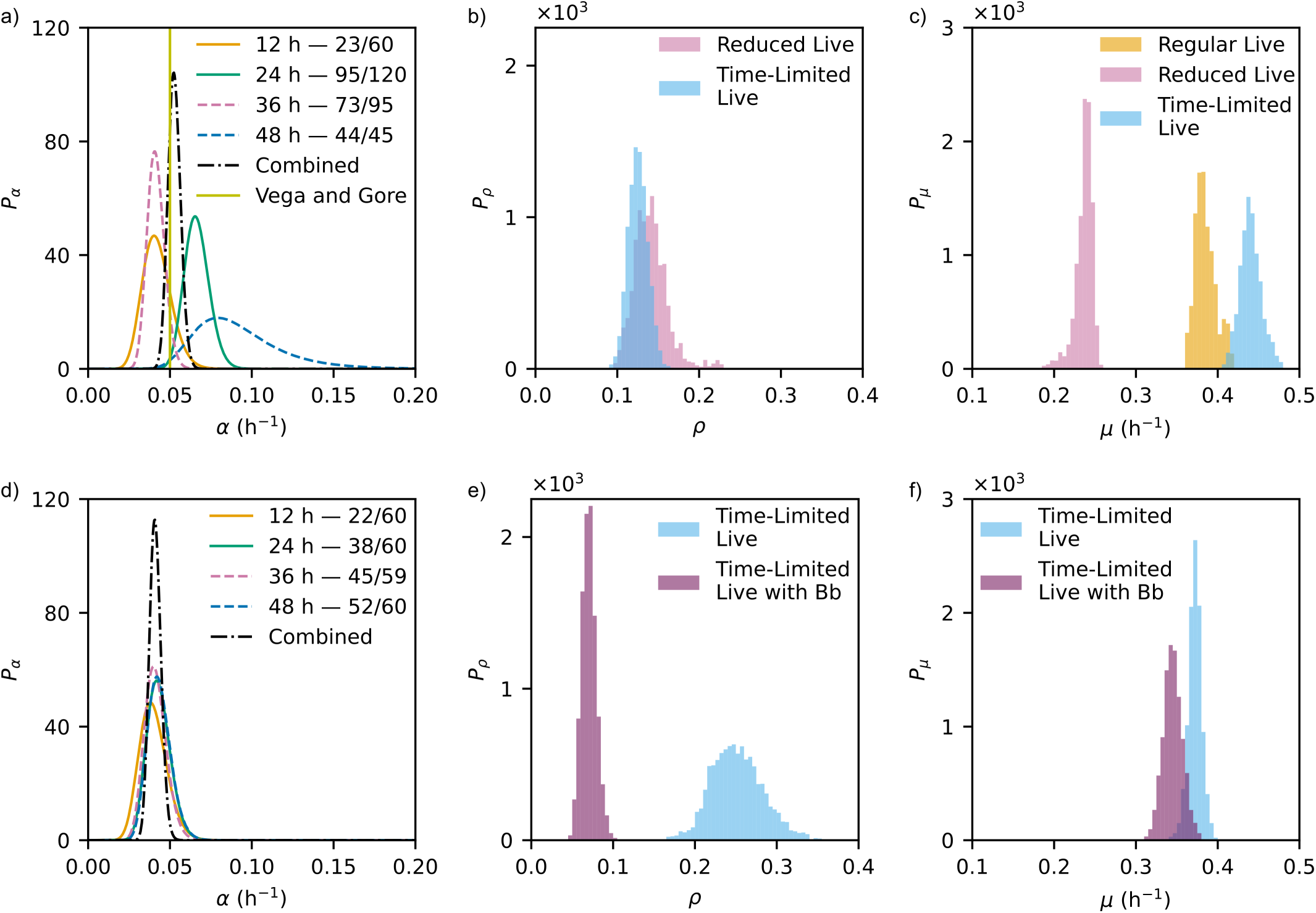
Inference of maximal colonization rate, environmental colonization scaling, and effective replication rate across feeding and predation conditions. a) Maximal colonization rate *α* under fully viable feeding conditions determined from the binary colonization experiment in S medium at pH 5.8. The inferred rates for *α* were compared with estimates from Ref. [20]. b) and c) show inferred distributions of the environmental scaling factor for colonization *ρ*, reflecting the availability of colonization-competent bacteria compared to the fully viable condition, and the effective growth rate *µ*, respectively. The inference was performed (detailed in Sec. 4.2) using the feeding conditions explored in Fig. 2, with regular, reduced live, and time-limited feeding shown in orange, pink, and blue, respectively. The *ρ* value for the regular live feed was fixed to 1, defining the baseline colonization condition under full live bacterial exposure. d) Maximal colonization rate *α* determined from the binary colonization experiment in low-salinity S medium at pH 7.3. e) and f) show inferred distributions of *ρ* and *µ* for the predation experiments from Fig. 3. Control and Bb-treated conditions are shown in purple and blue, respectively. Results in a) and d) were inferred jointly from measurements at 12 h, 24 h, 36 h, and 48 h (black dot-dashed line).

In S medium (pH 5.8), both reduced live and time-limited live diets yielded similar values of *ρ* (0.13 [0.10–0.19] and 0.14 [0.11–0.19], respectively; Fig. 4b), consistent with the lower availability of colonization-competent bacteria relative to the full live diet (Fig. S4).

The reduction of *ρ* was even more pronounced under predation. In low-salinity, pH 7.3 conditions, the inferred *ρ*decreased from 0.25 [0.19–0.32] in the time-limited control to 0.07 [0.05–0.09] in the Bb-treated group (Fig. 4e), indicating a substantial reduction in the effective environmental reservoir of colonizing bacteria. This decrease is consistent with direct measurements of environmental *E. coli* abundance (Fig. S5c).

In contrast, the inferred effective growth rate *µ* was largely conserved across conditions. In high-salinity, low-pH environments, *µ* was similar between regular live and time-limited feeding conditions (0.38 h^−1^ [0.36–0.41] versus 0.44 h^−1^ [0.42–0.47]; Fig. 4c), with overlapping distributions. A similar consistency was observed across pH-adjusted conditions, with overlapping inferred *µ* distributions between all *E. coli*-only diets (Fig. 4f; Fig. S8c). An exception was observed in the reduced live condition in the original S medium, where *µ* was lower (0.24 h^−1^ [0.21–0.25]), potentially reflecting temporal irregularities in population growth (Fig. 2c; see Sec. 3).

The reduction of *ρ* alongside conservation of *µ* suggests that environmental perturbations–whether through altered feeding schedules or predation–primarily impact bacterial population dynamics through changes in environmental reseeding. This pattern is consistent with a mechanism by which Bb reduces bacterial populations by limiting colonization and recolonization, rather than by altering in-host replication or clearance.

## 3 Discussion

In this work, we explore how stochastic early colonization drives divergence in downstream bacterial loads across hosts until reseeding by the environment shows hints of homogenizing bacterial population load variability before the worms die.

On account of this, we focus on the environmental recolonization mediated by the transfer of bacteria between hosts, and demonstrate that this plays a dominant role in shaping host-to-host variability. We find, perhaps unexpectedly, that a perturbation (in this case a bacterial predator), while reducing the gut bacterial count across worms, acts not by in-gut predation but by suppressing the transmission highway from host to host.

These results place stochastic colonization within a broader framework of host coupling through the environment, in which bacterial dispersal between hosts regulates population-level dynamics. Recent work has shown that low levels of dispersal lead to probabilistic colonization and divergent trajectories across hosts, whereas increased connectivity through inter-host transmission promotes homogenization via repeated reseeding events [3].

To quantitatively distinguish the contributions of colonization, recolonization, and within-host growth to these population-level dynamics, we developed a simulation-based inference framework capable of estimating effective kinetic parameters from the observed bacterial load distributions through a multiscale population regime. These population ranges are inaccessible to other existing inference tools, which are suited to either small or large populations by relying on either master equation [38–40] or mass action models [41–43], respectively–but never interpolating between both.

Using this framework, we confirmed the profound impact of environmental reseeding and subsequent transmission of *E. coli* by worms, linking environmental dynamics to the structure of population variability. We found that lower inferred values of *ρ*, the environmental scaling factor of colonization, explained the broader bacterial load distributions observed under time-limited feeding conditions, which amplify the effects of early stochastic colonization, without significant changes in *µ*.

We further applied this approach to a predator–prey perturbation, showing that the bacterial predator Bb reduces bacterial loads primarily by suppressing recolonization between hosts rather than by enhancing clearance of established populations within the gut. These results suggest that modulation of environmental connectivity provides a distinct mechanism for controlling microbiota dynamics.

One limitation of the current study is that residual bacteria associated with the worm exterior may not be completely removed by the washing and bleaching procedures. Consequently, some fraction of the bacteria contributing to continued colonization under the time-limited live diet may arise from bacteria remaining physically attached to the surface of the worms rather than exclusively from expelled gut populations. However, this possibility does not substantially alter the central interpretation of our results. Whether bacteria persist as free environmental cells or remain transiently associated with hosts, both mechanisms contribute to effective host-to-host coupling through recolonization-competent bacterial reservoirs. Importantly, the marked reduction in the fraction of colonized worms over time under the time-limited live with Bb diet, relative to the time-limited live control diet, still indicates a substantial suppression of effective environmental recolonization pressure, consistent with the reduced inferred environmental scaling factor *ρ* under predation. The inferred parameters were also biologically consistent across most environmental conditions. In particular, the effective growth rate *µ* remained largely stable across *E. coli*-only diets despite substantial variation in environmental exposure. This is consistent with the expectation that in-host replication and loss processes operate within a relatively stable physiological regime, as the pH and salinity conditions remain within the tolerance range of both *E. coli* [51] and *C. elegans* [52].

Notable deviations from this trend may reflect limitations of the assumption that colonization and growth rates remain constant over time. In the reduced live diet, the lower concentration of colonization-competent bacteria is expected to increase the waiting time to the first successful colonization event. Because substantial bacterial replication does not begin until colonization is established, this delay effectively postpones the onset of exponential growth. If the true colonization rate increases later in life–for example, due to age-dependent physiological changes in *C. elegans* such as reduced grinder efficiency [53]–then a model constrained to use a single time-independent colonization rate may compensate by inferring a higher average colonization rate and a lower replication rate, *µ*. In this case, the inferred parameters should be interpreted as coarse-grained averages over time-varying host physiology and environmental conditions rather than fixed biological constants.

Importantly, the simulation-based inference framework presented here is not intrinsically limited to time-independent models. Since inference is performed through forward simulation (see Sec. 4.2), more complex formulations—including time-varying colonization or growth rates—can be incorporated naturally. The primary constraint is therefore not methodological, but experimental: resolving such dynamics would require higher temporal resolution or complementary measurements to constrain additional parameters.

A key limitation of the current study lies in the nature of the measurements. Gut bacterial counts are obtained through destructive sampling, precluding longitudinal observation within individual hosts. Alternative approaches such as microscopy are also constrained, as visible light can alter worm behavior or induce phototoxic effects [54, 55].

Future work could integrate independently measured host ingestion and expulsion rates, or extend this framework to controlled systems such as chemostats or microfluidic chambers, where environmental inputs can be directly manipulated. These approaches may help further disentangle the contributions of stochastic colonization, environmental reseeding, and within-host growth to host-associated microbial population dynamics.

## 4 Methods

### 4.1 Experimental methods

#### Animals and bacterial strains

*Caenorhabditis elegans* (strain AU37: genotype glp-4(bn2) I; sek-1(km4) X), a mutant strain with temperature-sensitive sterility and enhanced sensitivity to pathogens, and *Escherichia coli* strain OP50 were obtained from the Caenorhabditis Genetic Center (CGC). *Bdellovibrio bacteriovorus* 109J (15143, ATCC) was purchased from the American Type Culture Collection.

#### Chemicals and Reagents

Triton X-100 (X100-100, Sigma-Aldrich), magnesium sulfate (M2643-500G, Sigma-Aldrich), citric acid monohydrate (C7129-500G, Sigma-Aldrich), ethylenediaminetetraacetic acid disodium salt (EDTA) (798681-100G, Sigma-Aldrich), iron (II) sulfate heptahydrate (F8633-250G, Sigma-Aldrich), manganese (II) chloride tetrahydrate (221279-100G, Sigma-Aldrich), potassium citrate tribasic monohydrate (25107-1KG, Sigma-Aldrich), zinc sulfate heptahydrate (Z0251-100G, Sigma-Aldrich), copper (II) sulfate pentahydrate (C8027-500G, Sigma-Aldrich), potassium phosphate monobasic (P0662-500G, Sigma-Aldrich), potassium phosphate dibasic (P3786-500G, Sigma-Aldrich), sodium phosphate dibasic anhydrous (S7907-500G, Sigma-Aldrich), sucrose (S-9378-1KG, Sigma-Aldrich), and HEPES (H4034-500G, Sigma-Aldrich) were purchased from Sigma-Aldrich. Thermo Fisher Scientific Inc. supplied tryptone (BP1421500, Fisher BioReagents™), yeast extract (DF0127179, Gibco™ Bacto™), and calcium chloride dihydrate (C79-500, Fisher Chemicals). The peptone (J636-500G, VWR) was obtained from VWR International-AMRESCO, LLC, Radnor, PA, USA. The cholesterol 95% (A1147018, Thermo Scientific), granulated agar (214510, BD Difco), and gentamicin sulfate powder (400-100P or 400100P010, GeminiBio) were purchased from Fisher Scientific. Sodium chloride (C829L41, MP Biomedicals) was purchased from Thomas Scientific.

#### Buffers, media, and plates

For culturing and colony/plaque-forming-unit detection of *E. coli* and Bb, LB broth and agar plates, YSPC broth, and double-layer plates were utilized. LB broth, containing tryptone (10 g/L), sodium chloride (10 g/L), and yeast extract (5 g/L), and LB plates, containing the same components with the addition of agar (15 g/L), were prepared using reverse osmosis (RO) water and adjusted to a pH of 7.0. YPSC broth was made from 1 g/L yeast extract, 1 g/L peptone, 0.5 g/L anhydrous sodium acetate adjusted to pH of 7.6 and enriched with 2 mM CaCl_2_ and 1 mM MgSO_4_ added after autoclaving [56]. Double-layer plates were prepared by preparing 1% and 0.6% agar solutions of 5.96 g/L HEPES buffer at a pH adjusted to 7.6, which were enriched with 2 mM CaCl_2_ and 1 mM MgPO_4_ after autoclaving [56]. After pouring the top layer of agar, an overnight OP50 bacterial culture was spun down at 5000×g for 6 min, resuspended in 25 ml PBS, and added to the top layer for a final concentration of OD_600_ = 1. Using an automatic pipette, 5 mL of the top layer was poured into each plate and spread by tilting. All solutions were sterilized in an autoclave before use in experiments.

#### *C. elegans* culturing

Experiments featuring *C. elegans* used media, plates, and buffers, including S medium, NGM plates, and M9 worm buffer, as previously described in the Wormbook [57]. The worms were subcultured by cutting the NGM agar from the previous culture and transferring pieces to NGM plates with fresh OP50 lawns no less than once every two weeks in accordance with the Wormbook [57].

#### *E. coli* culturing

*E. coli* of the OP50 strain was cultured overnight in LB broth at 37^◦^C at 250 rpm on an orbital shaker. The concentration of this culture was measured by determining its turbidity via the optical density at a wavelength of 600 nm (OD_600_). The *E. coli* was then centrifuged (460×g for 10 min) and resuspended in S medium for use in experiments.

Heat-killed OP50 was prepared by incubating 5×*E. coli* suspension, generated by resuspending the bacterial pellet in one-fifth of the volume required to achieve an OD_600_ = 1.0, followed by incubation at 80^◦^C for 60 min in the relevant S medium solution. For the experiments shown in Fig. 2, the *E. coli* used had passed through the gut of *C. elegans* at least once, except for one repeat of the time-limited live feed, which used bacteria sourced from the original supplier stock (see Supplementary Note 1).

#### *B. bacteriovorus* culturing and preparation

To culture the Bb used in all the predation experiments, the equivalent of 1 ml of OD_600_ = 0.2 *B. bacteriovorus* was cultured with 10 ml of overnight culture of OP50 in 50 ml of YSPC media at 29^◦^C at 170 rpm on an orbital shaker for 2 days. After two days, the lysate was centrifuged for 10 minutes at 460×g for 10 min, the supernatant collected, and the process repeated again for a total of 2 centrifugation steps. Following the second round of centrifugation, the supernatant was filtered through a 0.45 *µ*m syringe filter to remove any remaining prey. The turbidity of this filtrate was then measured at 600 nm, and aliquots were made for worm feeding experiments and kept at 4^◦^C overnight. On the day of the experiment, the *B. bacteriovorus* aliquots were centrifuged (7197×g for 30 min) and resuspended in S medium for use in the worm feeding experiments at a final concentration of OD_600_ = 0.2.

#### Synchronization of worm cultures

To synchronize the ages of the *C. elegans* used in the experiments, reproductively viable nematodes grown on NGM plates, as described above, were collected using sterile RO water. The eggs were isolated by treating the worms with a 10% 5M NaOH and 20% bleach solution, followed by centrifugation at 2000 g for 1 min to pellet the eggs. After three washes with sterile water, the eggs were inoculated along the edges of an

NGM plate with an OP50 lawn (Fig. 1)a). The plates were incubated at 25^◦^C for three days to obtain age-synchronized, gonadless adults.

#### Preparation of bacteria-free worms

Synchronized adult worms were collected by washing the NGM plates with M9 buffer containing 0.1% Triton X-100 and centrifugation at 2000×g, followed by two washes with M9 buffer to remove external bacteria. To remove any remaining live bacteria, the worms were then incubated in S medium with 100*µ*g/mL gentamicin and OD_600_ = 1 heat-killed OP50 bacteria for 24 h (Fig. 1)a). Subsequently, the worms were washed three times with a buffer – M9 Triton X-100, M9, and M9, sequentially – with centrifugation at each step to remove remaining traces of the antibiotic or heat-killed bacteria.

#### Worm gut colonization

Bacteria-free worms were grown in a 24-well plate at 25^◦^C with 250 rpm shaking incubator at a concentration of 200–800 worms/well inside a 1 ml S medium. Worms were fed on a certain diet of bacteria and then refed 24, 72, 120, and 168 hours after the initial live feeding by replacing the feeding solution with fresh bacterial suspension. The diet was varied between experiments (Fig. 1)a). For the standard experiment, each feeding consisted of 900 *µ*L S medium and 100 *µ*L 10× live OP50 bacterial suspension. For the time-limited feeding experiment, the first feeding remained the same as the standard, followed by sequential washes with M9-Triton X-100, M9, and then again with M9 at the 24 h mark to remove external live bacteria. Then, each subsequent feeding consisted of 800 *µ*L S medium and 200 *µ*L 5× heat-killed OP50 bacterial suspension. For the reduced live feed experiment, each feeding consisted of 800 *µ*L S medium, 198 *µ*L 5× heat-killed OP50 bacterial suspension, and 2 *µ*L 5× live bacteria for a ratio of 1:99 live:heat-killed bacteria. For the bacterial predator experiments, the pH of the S medium was adjusted to 7.3 due to the sensitivity of Bb to pH. The experiments followed the above-described protocol, but with the replacement of 100 *µ*L S medium for 100 *µ*L of OD_600_ = 2 Bb suspended in S medium, for a final concentration of OD_600_ = 0.2 Bb in each well. This corresponds to around 37.3 million PFUs per well. Total media volume per well remained 1 mL.

#### Nematode crushing

We used colony-forming units (CFU) to quantify the number of *E. coli* in the nematode guts at multiple time points in the experiment. To do this, we follow a previously published protocol that involves mechanically homogenizing the worms using a handheld homogenizer (BT704, Geno Technology Inc.) in a 50 *µ*L microcentrifuge tube and plating their gut contents onto LB agar plates [20]. First, the worms are washed with M9 and 0.1% Triton X-100 and then twice in M9 worm buffer. The worms were then purged by a 30 min incubation with 1× heat-killed OP50 bacterial suspension. The worms are then washed with 4^◦^C M9 and 0.1% Triton X-100 and centrifuged for 2 min at 2000×g in a 4^◦^C pre-cooled centrifuge. The worms are then suspended in 1 mL 4^◦^C M9 buffer with bleach solution at a 1:1000 dilution. The worms were incubated in this solution at 4^◦^C for 10 min. The worms were then rinsed in M9 plus 0.1% Triton X-100 solution and then resuspended in M9 with 1% Triton X-100 solution before being transferred to a 35 mm Petri dish. Between each rinse, worms were spun down for 1 min to ensure maximal worm retention.

Individual worms were pipetted with approximately 20 *µ*L of their surrounding solution into individual 600 *µ*L microcentrifuge tubes and disrupted using a motorized pestle. The disrupted worms were diluted in an additional 180 *µ*L PBS. Serial dilutions of the homogenized worms were made with PBS. 90 *µ*L of the dilutions were plated on LB agar plates, which were incubated overnight at 37^◦^C. This gives a dilution schedule of 22, 222, 2222, 22222 for the experiments shown in Fig. 2. When enumerating CFUs across dilutions, we report the count from the smallest dilution for which we had fewer than 300 visible colonies [20, 34].

#### Experiments with high salinity, pH 7.3 S media

The high salt, pH 7.3 version of the S media used in the experiments from Fig. S3 was created by increasing the pH from 5.8 to 7.3, with 5 M NaOH. The media were subsequently filtered through a 0.22*µ*m filter system. This increases the osmolarity of the S medium from ∼350 mOsm/L to ∼420 mOsm/L.

#### Experiments with low salinity, pH 7.3 S media for showing effects of predation on gut microbiota

For the experiments shown in Fig. 3, we adjusted the osmolarity to the tolerable range for Bb; the low salinity S medium was prepared by decreasing the base NaCl concentration from 100 mM to 10 mM. Then the pH adjustment was applied, and the media were subsequently filtered through a 0.22*µ*m filter system. This results in a final osmolarity of ∼220 mOsm/L for the predation-supporting medium.

Additionally, for these predation experiments and their controls, worms were resuspended and picked in M9 buffer without Triton X-100 to avoid detergent-induced toxicity to Bb [58]. These experiments also feature a 2.2 dilution, which is plated directly from the diluted homogenate. One exception occurred for Day 1 of a single Bb-treatment replicate, in which the diluted homogenate was not plated directly; these data were therefore excluded from calculations of the colonized worm percentage. After incubation, colony-forming units (CFUs) were enumerated for each plate. For Bb-treated worms, 90 *µ*L of the diluted homogenate from each individual worm was additionally plated on double-layer agar plates for plaque-forming unit (PFU) enumeration of Bb.

#### Environmental CFU quantification

On each day of nematode crushing, environmental bacterial counts from the wells were obtained by first carefully pipetting the environment to a separate microcentrifuge tube after the worms had settled. From this bacterial suspension, triplicate serial dilutions spanning 10^−3^ − 10^−7^ were produced, and 90 *µ*L of the appropriate dilutions were plated on LB plates to obtain CFUs.

#### Binary colonization rate experiments

For the binary experiments used to calculate baseline colonization rate described in Sec. 2.3, age-synchronized worms were prepared as described above and fed the standard diet of OD_600_ = 1 live OP50 for 12 h, 24 h, 36 h, or 48 h. Subsequently, the worms were washed and treated as in the worm gut colonization experiments before homogenizing. The homogenates were diluted with an additional 80 *µ*L of PBS and then plated in their entirety on an LB plate.

#### Bb predation in different environments

For the experiments of Bb predation inside different S media formulations (Fig. S5), we prepared Bb filtrates as described in the Bb culturing and preparation section. These filtrates were centrifuged at 7197×g for 30 min to pellet the Bb. A 24-well plate was prepared, each well containing OD_600_ = 1 live OP50 and OD_600_ = 0.2 Bb inside 1 mL of the chosen media. The plate was incubated at 25^◦^C inside a shaking incubator at 250 rpm. At different time points, the contents of 2-3 wells were collected in a cuvette before the OD_600_ was measured. All samples were zeroed with the corresponding buffer.

### 4.2 Theoretical inference methods

Our goal is to infer the dynamics governing bacterial population growth within the host. To this end, we quantify bacterial populations in individual worms over time.

The challenge is that these data are observed only indirectly in two distinct ways. First, bacterial populations within individual worms cannot be measured directly: CFU counts obtained through serial dilution and plating provide only a partial and stochastic readout of the true underlying population. Second, only a small subset of worms can be sampled at each time point. As a result, we do not observe longitudinal trajectories for individual worms, but only cross-sectional snapshots from different individuals at successive times.

To address the first source of uncertainty, we use REPOP, a method we previously developed to account for the stochasticity introduced by dilution and plating [34]. To address the second, we introduce a method that marginalizes over the unobserved within-host trajectories between measurements. Specifically, we model the latent distribution of bacterial population trajectories and use it to connect effective, nonlinear colonization and growth dynamics inside each worm to the experimentally observed CFU counts.

The goal of kinetic inference is therefore to estimate the parameters summarized in Fig. 1c. We model within-host bacterial population dynamics using the parameters (*α, K, ν, ω*), where *α* is the colonization rate (adhesion of *E. coli* from the environment to the gut), *ν* is the replication rate within the gut, *ω* is the loss rate due to expulsion or death, and *K* is the per-worm carrying capacity.

The following subsections address each component of this inference procedure. The next subsection focuses on the determination of the maximal colonization rate *α* from independent calibration experiments (Fig. 4a,d). Later, in experiments where the environmental bacterial population is perturbed (time-limited exposure or reduced viability; Fig. 2a and Fig. 3a), we represent the scaled colonization rate as *ρα*, where *ρ* denotes the fraction of bacteria competent to colonize the gut.

The subsequent subsection is dedicated to estimating the carrying capacity *K* from separate plating measurements. The final subsection presents the joint inference of the effective growth parameter *µ* and the colonization scaling factor *ρ* within a Bayesian framework that accounts for the indirect and stochastic nature of the observations.

#### Precalibrating the maximal colonization rate *α*

To precalibrate the maximal colonization rate *α*, we consider experiments in which entire worms are plated, such that colonization can be treated as a binary outcome (no colonies versus at least one colony). Under this measurement scheme, each observation reports whether at least one successful colonization event has occurred within an exposure time *t_i_*.

We model colonization as a memoryless process with rate *α*, such that the waiting time to the first colonization event is exponentially distributed. Accordingly, for each worm *i*, the probability of no colonization by time *t_i_* is 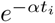, while the probability of at least one colonization event is 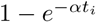.

We denote by *y_i_* ∈ {0, 1} the observed outcome for worm *i*, where *y_i_* = 1 indicates successful colonization and *y_i_* = 0 indicates no colonization. Therefore, *y_i_* follows a Bernoulli distribution with success probability 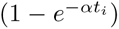, enabling direct estimation of *α* from the observed fraction of colonized worms across exposure times. As such, the likelihood of the observed data {*y_i_*} given *α* is thus

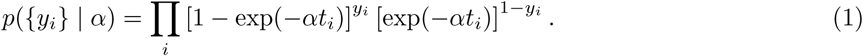

The experimental procedure is described in Methods (Binary colonization rate experiments for determining the maximal colonization rate). We estimate *α* within a Bayesian framework using a uniform prior, such that the posterior is proportional to the likelihood,

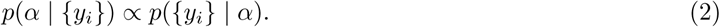

We report the maximum a posteriori estimate, 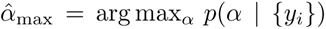, corresponding to the peak (black curve) in Fig. 2a and Fig. 3a under the respective experimental conditions.

#### Precalibrating the gut capacity *K*

In order to precalibrate the effective parameter *K*, we explicitly account for the substantial heterogeneity in end-stage bacterial loads observed both in our data and in prior studies [31]. Rather than assuming a single carrying capacity shared across worms, we model *K* as a worm-specific latent variable drawn from a distribution of ultimate carrying capacities.

We assume that, by the final day of regular feeding, most worms have reached an effective steady-state bacterial load under the experimental conditions. This is supported empirically by the observation that reconstructed bacterial counts plateau at similar orders of magnitude at late times rather than increasing indefinitely. We emphasize that this capacity is defined relative to the adult, end-of-life diet considered here, rather than as an absolute physiological ceiling.

We infer the distribution of *K* by reconstructing the final-day (day 9) bacterial count distribution of regularly fed worms in a modified S medium under conditions optimized for *E. coli* growth (pH 7.3) and non-stressed osmotic conditions for *C. elegans* ([*NaCl*] ≈ 100 mM), using REPOP [34]. REPOP accounts for the stochasticity introduced by dilution and plating to recover the underlying distribution of within-worm bacterial loads, represented as a mixture of Gaussians.

To exclude worms that were colonized late and may not have reached capacity by day 9, we retain only reconstructed populations exceeding 1000 bacteria and approximate the resulting distribution using the four rightmost Gaussian components. Stratified samples drawn from this distribution are then used as realizations of individual worm capacities in forward simulations. The resulting distribution is shown in Fig. S7.

#### Bayesian inference procedure for *µ* and *ρ*

We infer the effective growth rate *µ* and colonization scaling factor *ρ* from CFU data at discrete time points within a Bayesian framework that accounts for the indirect and stochastic nature of the measurements.

We assume that the underlying kinetic rates governing colonization, replication, and loss are effectively constant over the duration of the experiment. This is supported by the use of age-synchronized worms that have reached full adulthood prior to the onset of feeding, minimizing developmental changes that could otherwise alter gut physiology or feeding behavior on experimental timescales. While biological factors such as age-dependent changes in grinder efficiency may introduce weak time dependence in colonization dynamics, we do not model these effects explicitly. Instead, any residual temporal variation is absorbed into the inferred effective parameters, and *µ* and *ρ* should be interpreted as time-averaged quantities over the experimental window.

We further assume that ingested bacteria contribute to colonization proportionally to their viability. Under the experimental conditions considered here, worms are exposed to a fixed bacterial load that rapidly saturates feeding, such that the total number of bacteria entering the gut is effectively constant across conditions. In this regime, differences in colonization are driven primarily by the fraction of viable bacteria in the environment. We therefore model the scaled colonization rate as *ρα*, where *ρ* ∈ [0, 1] captures the fraction of bacteria competent to colonize relative to the fully viable condition.

Given these assumptions, we perform inference using a stochastic forward model of within-host dynamics, coupled to the measurement model describing dilution and plating. Posterior distributions over *µ* and *ρ* are obtained by comparing simulated observations to experimental CFU data, thereby capturing both intrinsic stochasticity and host-level variability.

Similarly, when introducing Bb, we treat predation effects as perturbations to baseline host–*E. coli* kinetics rather than explicitly modeling predator abundance. We expect external predation to reduce the environmental concentration of viable *E. coli*, thereby decreasing the pool of potential colonizers (represented in *ρ*).

#### Forward simulation

Simulations are performed using a hybrid stochastic algorithm to generate population trajectories {*n*(*t*; *α, µ, ρ*)}*_t_*_∈[0*,T* ]_, where *n*(*t*; *α, µ, ρ*) denotes the gut bacterial population size at time *t* under the specified kinetic parameters.

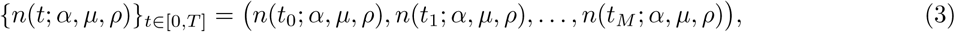

Each simulated trajectory additionally depends on a latent, worm-specific carrying capacity parameter *κ_i_*, representing the ultimate bacterial capacity of the host. For each realization, a value of *κ_i_* is drawn from the distribution of host capacities *K* mentioned in S7 and held fixed over the trajectory. The resulting population trajectories are therefore

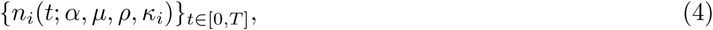

where (*α, µ, ρ*) denote the kinetic parameters and *κ_i_* is the sample fom the distribution obtained in S7.

To generate these trajectories, we simulate the underlying stochastic dynamics using a hybrid scheme combining exact stochastic simulation and approximate tau-leaping. At low population sizes, we use the direct Gillespie algorithm [59] to accurately capture discrete stochastic events. As the population increases and reaction events become frequent, exact simulation becomes computationally prohibitive. We therefore switch to an approximate simulation scheme once the population exceeds a threshold (here *n >* 250), in which multiple reaction events are evolved collectively over short time intervals under the assumption that reaction rates remain approximately constant. At these large populations, the time step is chosen adaptively such that the expected change in population over each interval remains below 1% of the current population size. This approximate scheme, in which multiple reaction events are evolved collectively over short intervals, is commonly referred to as tau-leaping [60] and preserves accuracy in the low-copy-number regime while enabling efficient simulation at large population sizes.

In these simulations, we account for changes in environmental viability by modifying the colonization rate. Under fully viable feeding conditions, colonization occurs at its maximal rate *α*. In conditions where bacterial viability is reduced, the effective colonization rate is scaled as

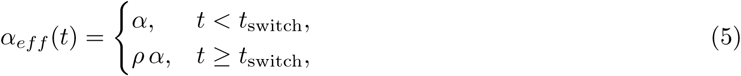

where *ρ* ∈ [0, 1] represents the fraction of bacteria capable of successful colonization, and *t*_switch_ the time at which the environment change happens. In full live feeding experiments, *ρ* = 1, such that *α_eff_* (*t*) = *α* at all times.

Full code for the simulations is available in our GitHub repository [61]

#### Learning the kinetics from forward simulation

Our goal is to infer the effective parameters *µ* and *ρ* from CFU measurements. For each worm *i*, we observe a colony count *c_i_* at dilution factor *ϕ_i_* and exposure time *t_i_*. Following the protocol established in REPOP [34], we identify the first dilution in a predefined series that falls below a detection threshold (see Sec. 4.1) with associated time *t_i_*. This induces a stochastic relationship between the observed pair (*c_i_, ϕ_i_*) and the underlying, unobserved within-host bacterial population size *n*(*t_i_*).

In probabilistic language and denoting the dataset by {(*c_i_, ϕ_i_*)} = {(*c_i_, ϕ_i_*)}*_i_*_∈1:*I*_, our goal is to compute

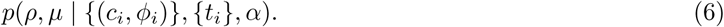

By Bayes’ theorem, the distribution above, termed the posterior distribution over parameters, is given by

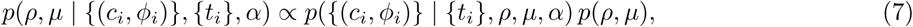

where *p*(*ρ, µ*) denotes the prior over the effective growth and colonization parameters, and *p*({(*c_i_, ϕ_i_*)} | {*t_i_*}*, ρ, µ, α*) is the likelihood of the observed CFU data under the model.

Assuming conditional independence across worms, the likelihood factorizes as

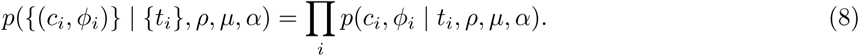

Each term *p*(*c_i_, ϕ_i_* | *t_i_, ρ, µ, α*) represents the probability of the observed CFU outcome for worm *i* and is obtained by marginalizing over the latent within-host population trajectories and the dilution–plating measurement process.

More specifically, for each worm *i* observed at time point *t_i_* we marginalize over the latent within-host bacterial population size *n_i_* to determine the probability *p*(*c_i_, ϕ_i_*|*θ, t_i_*) of observing a CFU count *c_i_* for a dilution factor *ϕ_i_* given parameters *θ*.

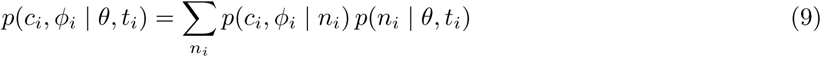

Noting that per-worm capacity *κ_i_* is a latent host trait and *p*(*κ_i_*) is the precalibrated capacity distribution described in the assumptions, we can express the likelihood of any single realization of (count, dilution) datapoint in a worm as

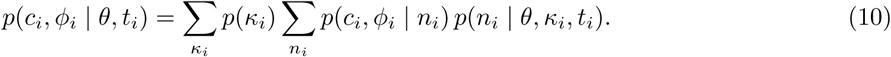

Equation 10 defines the conceptual likelihood after marginalizing over host-level capacity heterogeneity; in practice, this marginalization is performed implicitly through Monte Carlo simulation. We can calculate the sampling likelihood *p*(*c_i_, ϕ_i_*|*n_i_*) for bacterial number *n_i_* dilution *ϕ* and CFU number *c_i_*, as described in our previous work [34]. However, calculating *p*(*n_i_*|*θ, κ_i_, t_i_*) via a chemical master equation would be computationally prohibitive, thus we use a Monte Carlo approximation. For a given parameter set *θ*, we generate *S* = 2^20^ independent simulated trajectories using the forward model described in Sec. 4.2, yielding samples 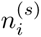. The likelihood is then approximated as

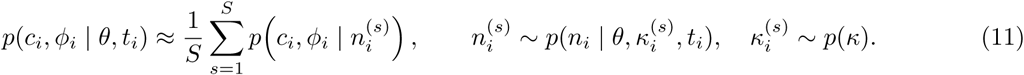

With the likelihood in (11), the factorized likelihood in (8), and Bayes’ theorem in (7), the posterior over (*ρ, µ*) is, in principle, fully specified. In practice, however, evaluating this posterior is computationally demanding because each likelihood term requires stochastic forward simulation and Monte Carlo averaging. We therefore proceed in two stages: first, a grid-based evaluation to identify the global structure of the posterior landscape; and second, Markov chain Monte Carlo (MCMC) sampling to more efficiently characterize the posterior distribution.

#### Grid-based likelihood evaluation and Markov chain Monte Carlo sampling for *ρ* and *µ*

Inference reduces to a two-dimensional parameter space over (*ρ, µ*). We restrict this space to biologically plausible bounds, with *ρ* ∈ (0, 1], since *ρ* is a scaling factor on the maximal colonization rate, and *µ* ∈ (0, 3], where the upper bound is motivated by reported maximal replication rates of *E. coli* in rich media [62]. In practice, we find that for all experiments the posterior mass is well contained within *ρ* ≤ 0.5 and *µ* ≤ 0.5, indicating that these bounds are conservative.

We first evaluate the likelihood over a grid in (*ρ, µ*) with resolution 0.02 in each dimension to obtain a coarse-grained view of the posterior landscape and identify regions of high posterior support for each experiment (Fig. S9). Because the likelihood is estimated via stochastic forward simulation, it is subject to Monte Carlo variability arising from finite sampling of latent trajectories. Consequently, the highest-scoring point on the grid may be sensitive to simulation noise and does not necessarily provide a stable point estimate.

To obtain a more robust characterization of posterior uncertainty, we use the high-posterior regions identified by the grid search to initialize MCMC sampling. Specifically, we define a subregion of the parameter space by retaining grid points whose posterior value is within a factor of *e*^20^ (i.e., within 20 in log-posterior) of the maximum. This yields a bounded region [*µ*_low_*, µ*_high_] × [*ρ*_low_*, ρ*_high_], and we initialize the MCMC chain at the midpoint of this region.

We then employ a Metropolis–Hastings algorithm with Gaussian random-walk proposals, where the proposal widths are set to 0.025 (*µ*_high_ − *µ*_low_) and 0.025 (*ρ*_high_ − *ρ*_low_) in each respective dimension. By repeatedly evaluating nearby parameter values, this procedure reduces sensitivity to local Monte Carlo fluctuations and yields samples from the posterior distribution over (*ρ, µ*). We collect 10^4^ samples per experiment, leading to the posterior samples shown in Fig. 4b, c, e, f, as well as Fig. S8b, c. Full code for the calculations is available in our GitHub (see Code availability).

## Data availability

Plate images are available on https://zenodo.org/records/20128602 (DOI: 10.5281/zenodo.20128602) and figure data is available https://github.com/PessoaP/Celegans_inference.

## Code availability

Code for the simulation-based inference is available at https://github.com/PessoaP/Celegans_inference.

## Acknowledgments

Figs. 1, 2, 3, as well as Figs. S2, S3 were created using BioRender assets. Tashev, S. (2026) https://BioRender.com/i81vmg5. The authors acknowledge Research Computing at Arizona State University for providing HPC resources that have contributed to the research results reported within this paper [63].

