## Supplemental Information for "Stochastic colonization and host-to-host transmission shape gut bacterial variability"

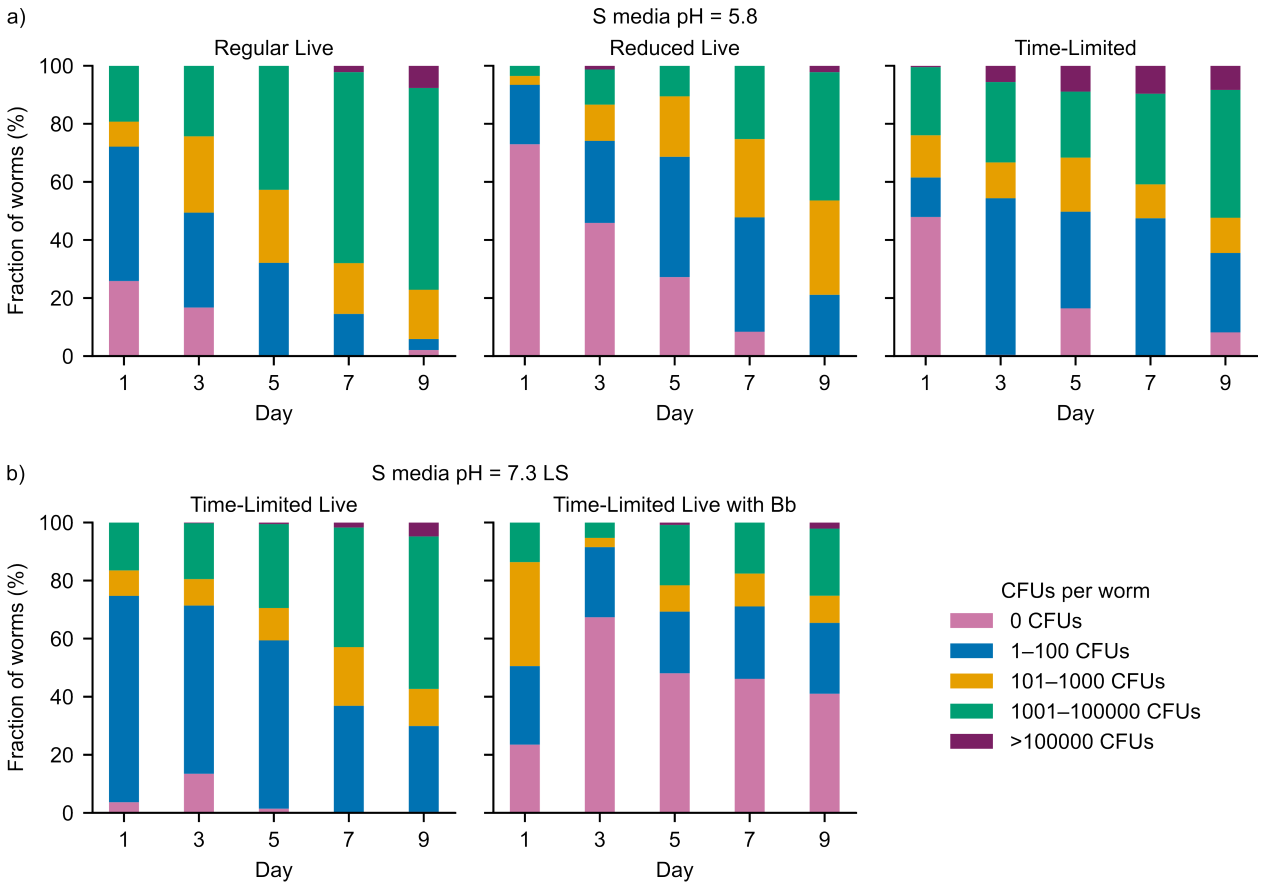


**Figure S1: Diet, environment, and predation shape the temporal trajectory of bacterial load distributions.** Breakdown of the probability distributions of REPOP-reconstructed bacterial counts within the gut of worms under different feeding conditions: a) S medium pH = 5.8, b) S medium pH = 7.3 with low salinity (LS). REPOP reconstructed bacteria counts are binned to 0 CFUs, 1-100 CFUs, 101-1000 CFUs, 1001-100000 CFUs, and $\geq$100000 CFUs.

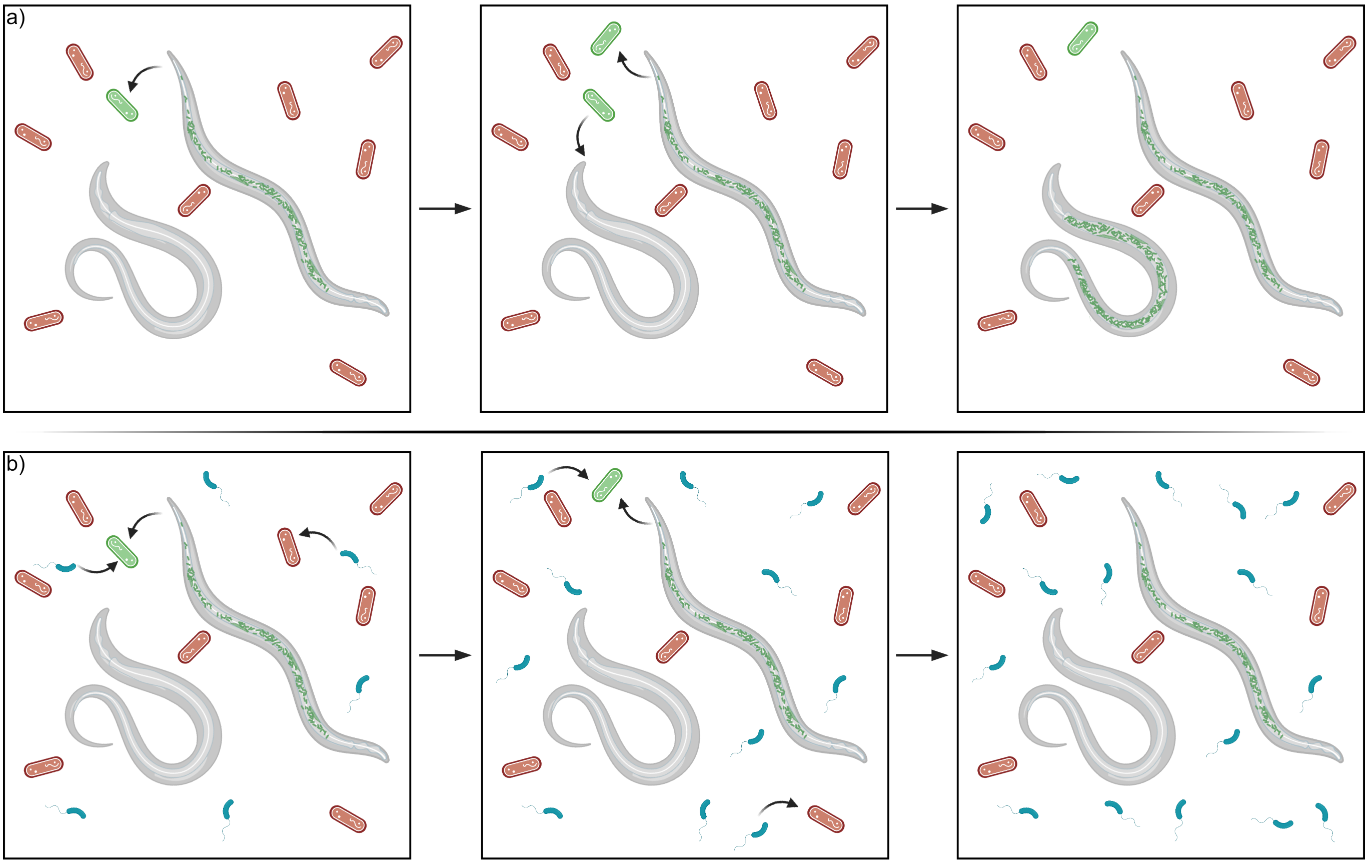


**Figure S2: Illustration of how recolonization arises from environmental regrowth of expelled live gut bacteria, but is suppressed by predation.** a) Illustration showing how the environment contains live bacteria (green) even if only a heat-killed (orange) diet is fed, due to the excretion from colonized worms. b) This reseeding effect is greatly reduced by Bb (blue), which feeds on both the excreted live bacteria (green), as well as the heat-killed bacteria (orange).

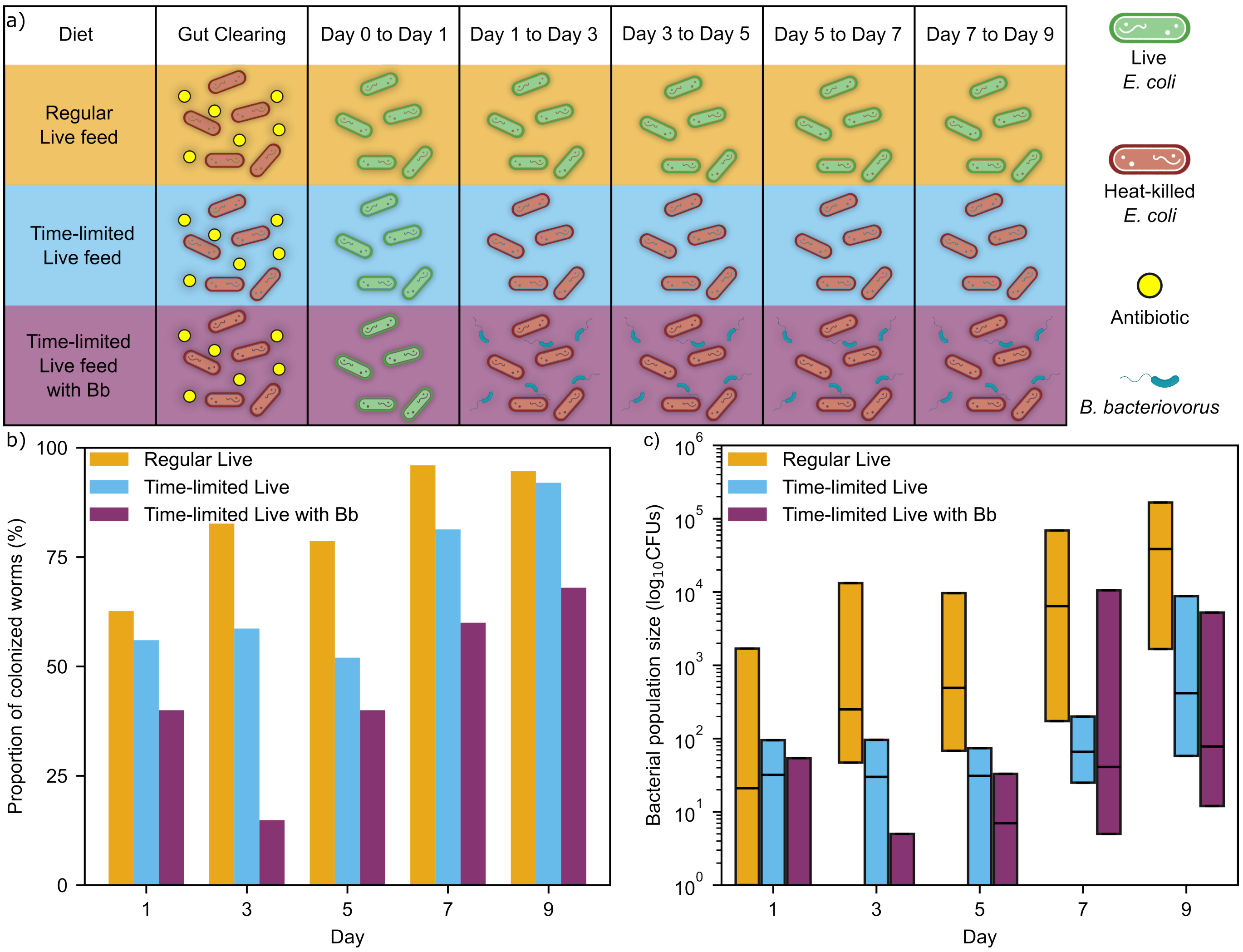


**Figure S3: Under higher pH, gut bacterial loads from regular live diet are elevated, yet broad population variability limits detectable predation effects.** a) Nematode diet plans featuring different periods of live (green) and heat-killed (red) bacteria, after an initial day of gut clearing with an antibiotic (yellow). Experiments were conduced in S medium of pH 7.3, higher than the standard S medium of pH 5.8. b) Mean percentage of colonized worms from the regular, time-limited, and time-limited live fiet with *B. bacteriovorus* (Bb) in orange, blue, and purple, respectively. The day labels represent the time point at which the worms were sampled. Sampled worms were not refed with bacteria on the day of crushing. c) Box plot of the quartiles of the gut bacterial population reconstructed by REPOP with the regular, time-limited feed, and the time-limited feed with Bb in orange, blue, and purple, respectively.

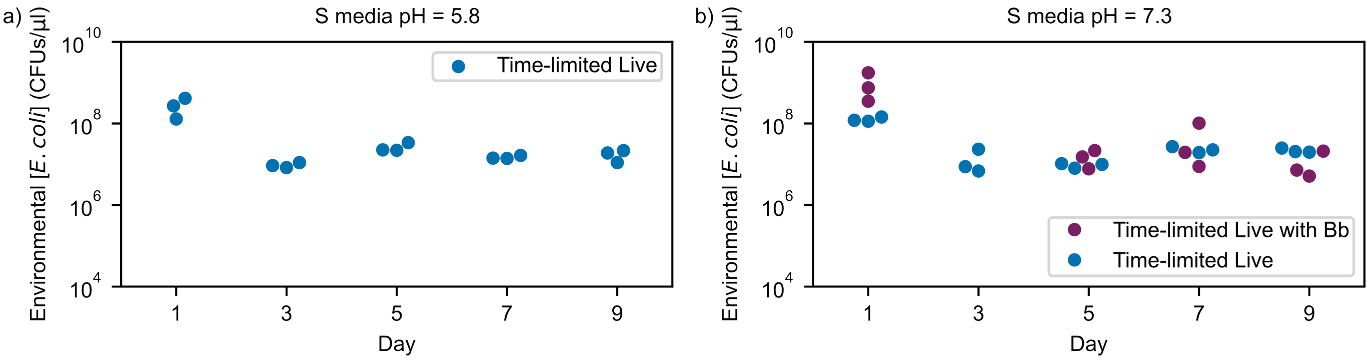


**Figure S4: Environmental bacterial concentration over time remains identical across all the high salinity time-limited live fiets, regardless of pH or predation.** Reconstructed environmental bacterial counts at different time points inside the wells of the time-limited feed without Bb (blue) and with Bb (purple) in the high salinity S media at a) pH = 5.8 and b) pH = 7.3.

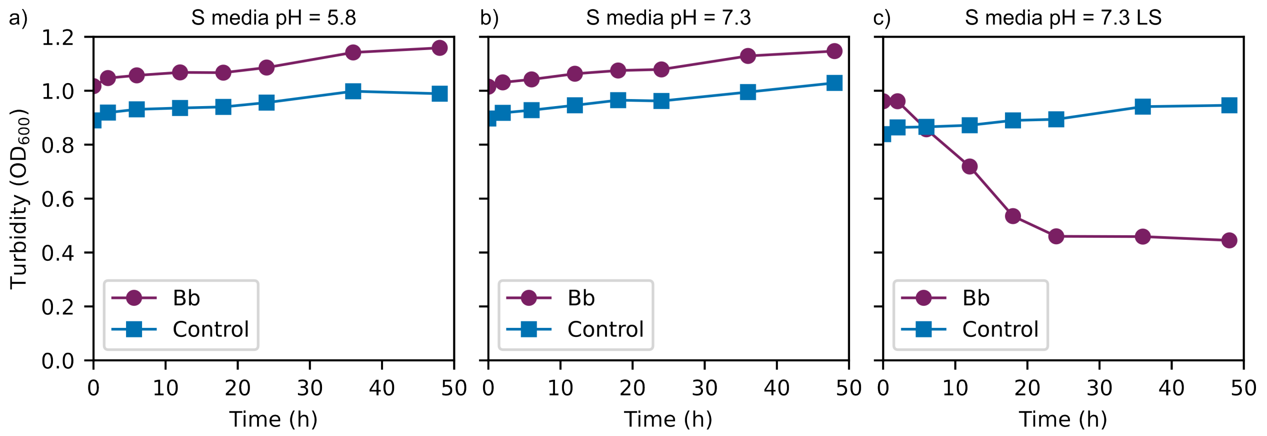


**Figure S5: Bb lowers optical density measurements only in low-salinity, high-pH medium.** Turbidity measurements ($OD_{600}$) of Bb-*E. coli* co-culture (purple) and control containing only *E. coli* (blue) over time. The bacterial concentrations were the same as in the nematode feeding experiments, OP50 at $OD_{600} = 1$ and Bb at $OD_{600} = 0.2$. a) S medium at pH = 5.8, b) S medium at pH = 7.3, c) low salinity S medium at pH = 7.3. All Bb–*E. coli* co-culture conditions begin with higher $OD_{600}$ values than the control, consistent with increased total cellular biomass at initialization. LS- low salinity.

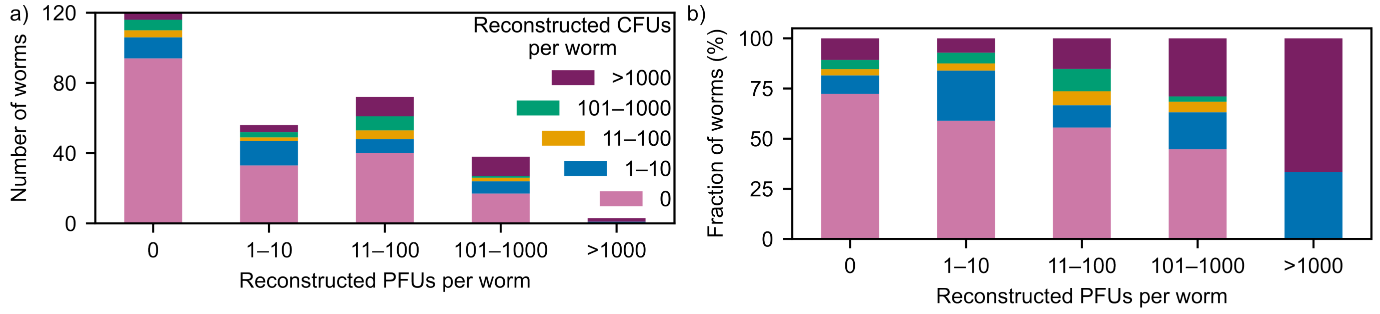


**Figure S6: *E. coli* population distributions across stratified Bb abundance levels within worms.** Stacked bar plot showing the quantities of CFUs (pink: 0, blue: 1 to 10, orange: 11 to 100, green: 101 to 1000 CFUs, purple: ≥1000 CFUs) in worms with different quantities of PFUs. a) shows the absolute number of worms that correspond to each Bb count strata, while b) is the relative fraction of worms in each strata. Conditioned on Bb abundance, the distribution of worms across *E. coli* population states remains relatively similar.

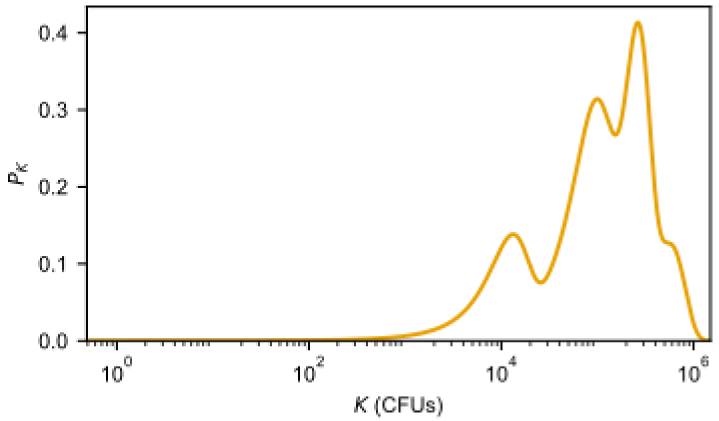


**Figure S7: Probability distribution of precalibrated worm capacities used in inference.** *E. coli* count distribution was reconstructed using REPOP from the final day (day 9) data of the high salinity, pH 7.3 regular live feeding condition. The four highest Gaussians in the inferred Gaussian mixture model are taken as the capacity distribution, with $K < 1000$ set to a probability of 0.

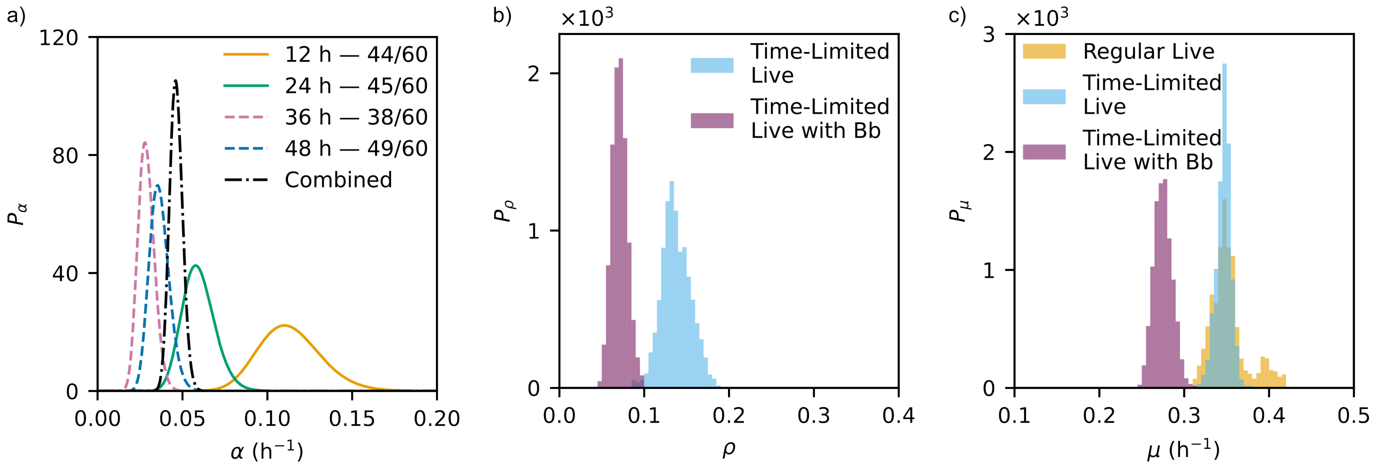


**Figure S8: Results from the stochastic inference pipeline for pH = 7.3 high-salinity condition.** a) Colonization rate $\alpha$ for high-salinity S medium with pH = 7.3 determined by the binary experiment, taken from 4 different time points (12 h, 24 h, 36 h, and 48 h), and combined (black dot-dashed line). b) and c) show the probability distributions of the environmental scaling factor for colonization $\rho$ and the effective replication $\mu$, respectively. The data for inference from the different diets explored in Fig S3 with regular live, time-limited live control, and time-limited live with Bb feed in orange, blue, and purple, respectively. The $\rho$ for the regular feed is set to 1, as that feed is considered the baseline colonization rate from the full live feed concentration.

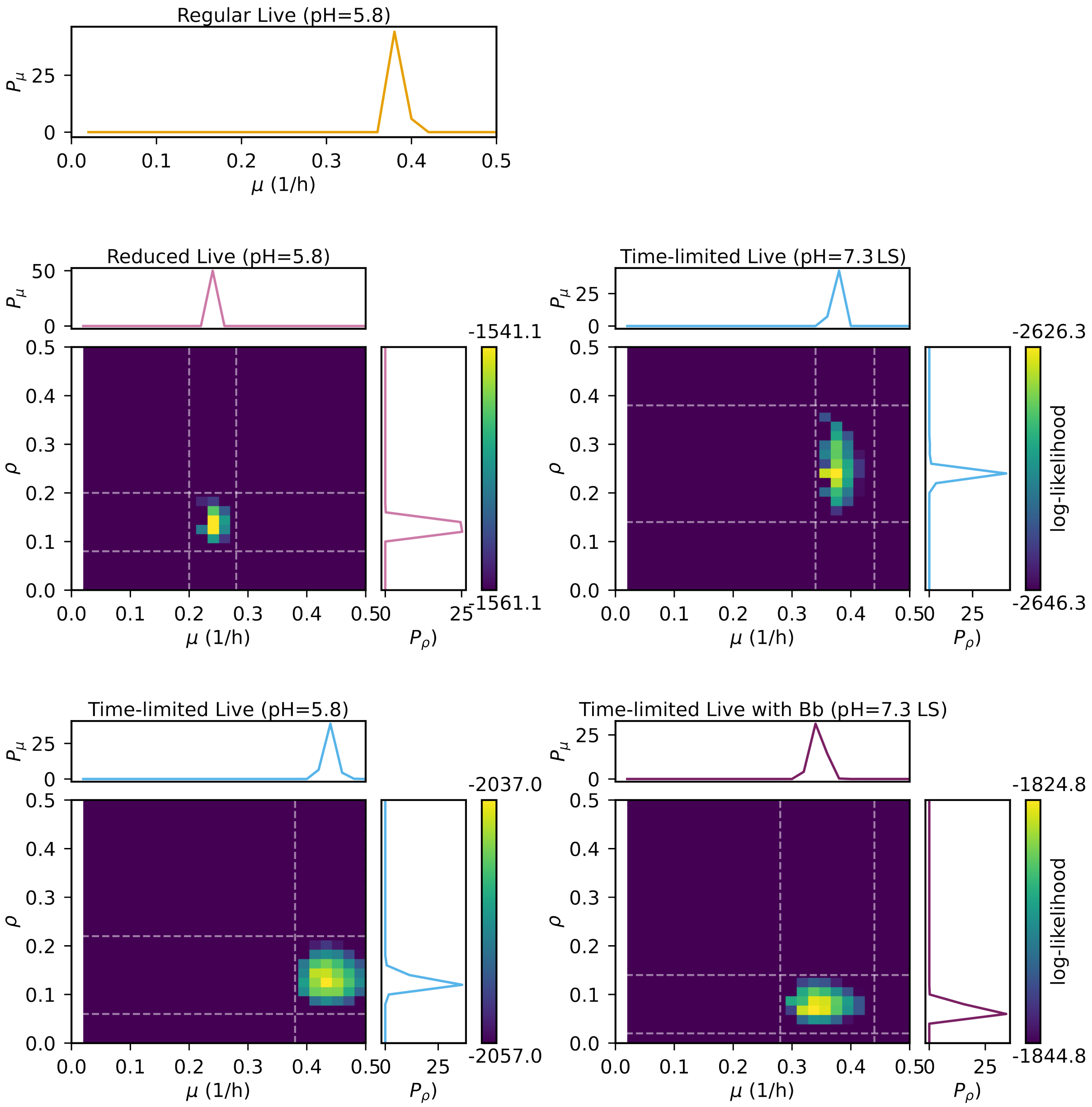


**Figure S9: Initial grid-based likelihood search for plausible values of $\mu$ and $\rho$.** A coarse grid search was performed over biologically plausible ranges of the effective replication rate $\mu$ and colonization scaling factor $\rho$. For regular live-feeding conditions, $\rho$ is fixed at 1, reflecting the assumption that the fraction of non-viable bacteria in the environment is negligible. The displayed region is restricted to $\rho \in [0, 0.5]$ and $\mu \in [0, 0.5]$ for visualization. The white dashed white lines delimit the region outside of which all computed likelihood values are smaller than the maximum by a factor of at least $e^{20}$. We use this region to initialize the MCMC, with step sizes chosen proportional to its widths, as described in Sec. 4.2: Grid-based likelihood evaluation and Markov chain Monte Carlo sampling for $\rho$ and $\mu$. LS- low salinity.

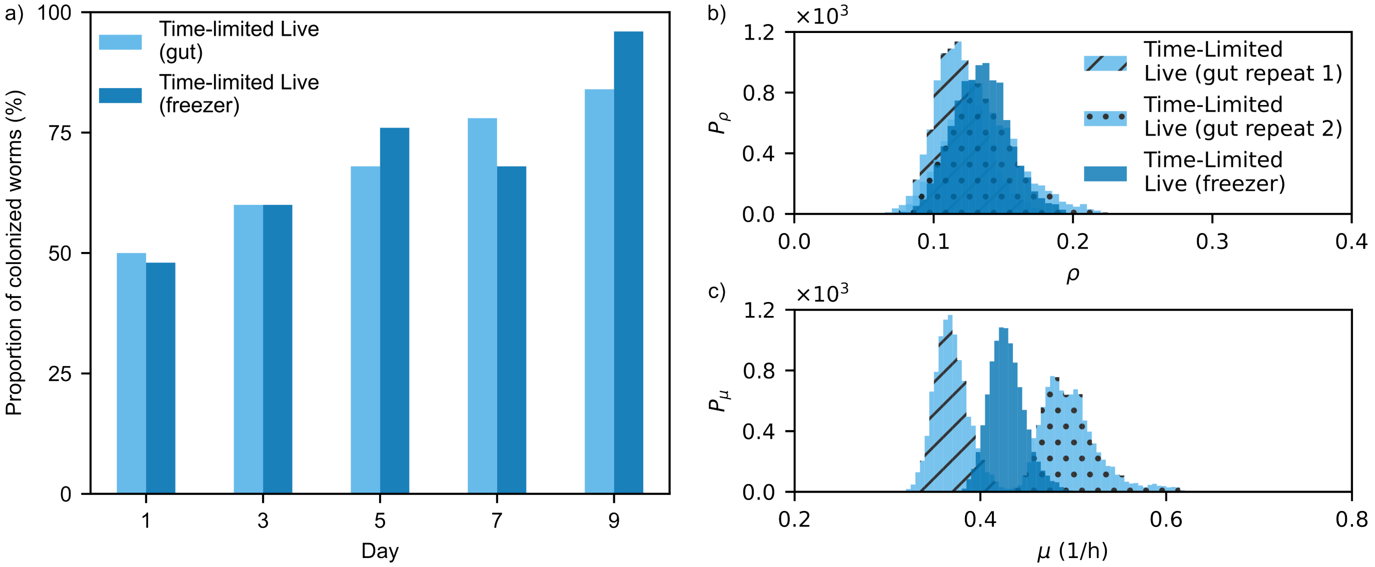


**Figure S10: Comparison between the microbiota derived from *E. coli* passed through the gut of *C. elegans* and those from a standard freezer stock.** a) Comparison of the colonization of worms with the gut stock (light blue) and the freezer stock (dark blue) when exposed to the time-limited live diet. b) and c) show the inferred rates of the environmental colonization factor $\rho$ and the replication rate $\mu$, respectively. The data from the gut *E. coli* stock repeats are in hatched and dotted light blue bars, while the freezer stock data is in dark blue bars. The gut stock results are broken down into the different experimental repeats.

**Supplemental Note 1.** The bacteria used in the experiments in the S media with pH 5.8, which are shown in Fig. 2, have been passed through the gut of *C. elegans* at least once, except for one repeat of the time-limited live diet, which has been done with a fresh stock of *E. coli* from the freezer. Comparing the percentage of colonized worms between the two repeats done with the gut stock and the repeat with the freezer bacterial stock, there are small differences in the percentage of colonized worms without any correlation trend observed between them (Fig. S10a). When it comes to the inference of the rates for the time-limited rates, the results for the $\rho$ values are similar, with the modes of the distribution around 0.12 (Fig. S10b). Moreover, the $\mu$ distribution for the freezer stock repeat has a center at around 0.43 $h^{-1}$, which is situated between the results for the two repeats of the freezer stock *E. coli* (Fig. S10c). Thus, we postulate that no significant genetic or behavioral difference was sustained from the passage of the bacteria through the nematode gut, and we have combined the freezer and the gut experiments in the analysis of Fig. 2 and Fig. 4, and we refrain from statistically comparing the results from these experiments directly to the rest of the experiments.

**Table S1:** Percentage of colonized worms across diets over time cultured in S medium at pH = 5.8. Values of replicates are shown as percentages.

| Day | Regular Live (replicates %) | Reduced Live (replicates %) | Time-limited Live (replicates %) |
|---|---|---|---|
| 1 | (60.0, 60.0, 56.0) | (36.0, 12.0, 20.0) | (48.0, 52.0, 48.0) |
| 3 | (88.0, 100.0, 32.0) | (36.0, 40.0, 60.0) | (64.0, 56.0, 60.0) |
| 5 | (96.0, 100.0) | (64.0, 64.0, 44.0) | (64.0, 72.0, 76.0) |
| 7 | (88.0, 100.0, 100.0) | (88.0, 92.0, 56.0) | (76.0, 80.0, 68.0) |
| 9 | (96.0, 100.0, 100.0) | (92.0, 92.0, 92.0) | (80.0, 88.0, 96.0) |

**Table S2:** Quartiles of the REPOP reconstructed populations throughout the different experiments conducted in S medium at pH = 5.8.

| Day | Regular Live (CFUs) | Reduced Live (CFUs) | Time-limited Live (CFUs) |
|---|---|---|---|
| 1 | (0, 36, 296) | (0, 0, 13) | (0, 24, 832) |
| 3 | (0, 13, 758) | (0, 23, 132) | (3, 73, 6816) |
| 5 | (49, 346, 6903) | (0, 15, 153) | (17, 106, 15857) |
| 7 | (239, 9705, 21960) | (16, 111, 2538) | (7, 127, 22273) |
| 9 | (1211, 11190, 25609) | (120, 377, 7330) | (47, 1382, 27477) |

**Table S3:** Percentage of colonized worms across diets over time cultured in S medium at pH = 7.3. Values of replicates are shown as percentages.

| Day | Regular Live (replicates %) | Time-limited Live (replicates %) | Time-limited Live with Bb (replicates %) |
|---|---|---|---|
| 1 | (72.0, 72.0, 44.0) | (56.0, 60.0, 52.0) | (60.0, 32.0, 28.0) |
| 3 | (80.0, 100.0, 68.0) | (48.0, 52.0, 76.0) | (28.0, 8.3, 8.0) |
| 5 | (88.0, 80.0, 68.0) | (44.0, 48.0, 64.0) | (56.0, 20.0, 44.0) |
| 7 | (96.0, 92.0, 100.0) | (80.0, 92.0, 72.0) | (56.0, 72.0, 52.0) |
| 9 | (92.0, 96.0, 96.0) | (84.0, 96.0, 96.0) | (56.0, 76.0, 72.0) |

**Table S4:** Quartiles of the REPOP reconstructed populations throughout the different experiments conducted in S medium at pH = 7.3

| Day | Regular Live (CFUs) | Time-limited Live (CFUs) | BB Time-limited Live (CFUs) |
|---|---|---|---|
| 1 | (1, 21, 1691) | (0, 32, 95) | (0, 0, 54) |
| 3 | (47, 250, 13200) | (0, 30, 96) | (0, 0, 5) |
| 5 | (68, 492, 9655) | (0, 31, 74) | (0, 7, 33) |
| 7 | (173, 6414, 69164) | (25, 66, 200) | (5, 41, 10544) |
| 9 | (1670, 38583, 166411) | (58, 416, 8792) | (12, 78, 5251) |

**Table S5:** Percentage of colonized worms across diets over time cultured in the low salinity S medium at pH = 7.3. Values shown as replicate percentages. One of the repeats is removed from the plots due to an uneven dilution (see Sec. 4.1).

| Day | Time-limited Live (replicates %) | Time-limited Live with Bb (replicates %) |
|---|---|---|
| 1 | 72.0, 88.0, 88.0 | 48.0, 70.83, (12.0) |
| 3 | 96.0, 88.0, 54.17 | 48.0, 4.0, 28.0 |
| 5 | 96.0, 100.0, 91.67 | 36.0, 20.83, 60.0 |
| 7 | 100.0, 100.0, 100.0 | 44.0, 32.0, 36.0 |
| 9 | 92.0, 100.0, 100.0 | 40.0, 40.0, 72.0 |

**Table S6:** Quartiles of the REPOP reconstructed populations throughout the different experiments conducted in S medium at pH = 7.3 with the low salinity.

| Day | Time-limited Live (CFUs) | BB Time-limited Live (CFUs) |
|---|---|---|
| 1 | (2, 13, 112) | (1, 66, 651) |
| 3 | (1, 18, 146) | (0, 0, 2) |
| 5 | (8, 30, 3915) | (0, 1, 514) |
| 7 | (43, 373, 7096) | (0, 1, 231) |
| 9 | (57, 1452, 12283) | (0, 4, 1047) |

**Table S7:** Median colonization rates $\alpha$ across different S medium variations.

| Medium | Median $\alpha$ (1/h) | 95% CI (1/h) |
|---|---|---|
| S medium pH = 5.8 | 0.05232 | [0.04522, 0.06029] |
| S medium pH = 7.3 | 0.04595 | [0.03895, 0.0.05388] |
| S medium pH = 7.3 low salinity | 0.0407 | [0.03419, 0.04812] |

**Table S8:** Inferred values for $\rho$ and $\mu$ values across S medium conditions and experiments. The median and the 95% CI are given with the 2.5th and 97.5th quantiles in brackets. LS- low salinity. For the Regular Live diet experiments, the $\rho$ is set to 1.

| S medium variant | Experiment | $\rho$ (median [95% CI]) | $\mu$ (median [95% CI], 1/h) |
|---|---|---|---|
| pH = 5.8 | Regular Live | 1.0000 [1.0000, 1.0000] | 0.3829 [0.3644, 0.4139] |
| pH = 5.8 | Reduced Live | 0.1393 [0.1068, 0.1945] | 0.2386 [0.2062, 0.2514] |
| pH = 5.8 | Time-limited Live | 0.1251 [0.1028, 0.1491] | 0.4394 [0.4158, 0.4692] |
| pH = 7.3 | Regular Live | 1.0000 [1.0000, 1.0000] | 0.3487 [0.3196, 0.4089] |
| pH = 7.3 | Time-limited Live | 0.1360 [0.1054, 0.1727] | 0.3473 [0.3273, 0.3624] |
| pH = 7.3 | Time-limited Live + Bb | 0.0708 [0.0542, 0.0904] | 0.2747 [0.2553, 0.2965] |
| pH = 7.3 LS | Time-limited Live | 0.2498 [0.1937, 0.3174] | 0.3724 [0.3552, 0.3889] |
| pH = 7.3 LS | Time-limited Live + Bb | 0.0702 [0.0545, 0.0895] | 0.3447 [0.3227, 0.3694] |
